# Structure of the vasopressin hormone-V2 receptor-β-arrestin1 ternary complex

**DOI:** 10.1101/2022.02.11.480047

**Authors:** Julien Bous, Aurélien Fouillen, Hélène Orcel, Stefano Trapani, Xiaojing Cong, Simon Fontanel, Julie Saint-Paul, Joséphine Lai-Kee-Him, Serge Urbach, Nathalie Sibille, Rémy Sounier, Sébastien Granier, Bernard Mouillac, Patrick Bron

## Abstract

Arrestins interact with G protein-coupled receptors (GPCRs) to stop G protein activation and to initiate key signaling pathways. Recent structural studies shed light on the molecular mechanisms involved in GPCR-arrestin coupling, but whether this process is conserved among GPCRs is poorly understood. Here, we report the cryo-electron microscopy active structure of the wild-type arginine-vasopressin V2 receptor (V2R) in complex with β-arrestin1. It reveals an atypical position of β-arrestin1 compared to previously described GPCR-arrestin assemblies, associated with an original V2R/β-arrestin1 interface involving all receptor intracellular loops. Phosphorylated sites of the V2R C-terminus are clearly identified and interact extensively with the β-arrestin1 N-lobe, in agreement with structural data obtained with chimeric or synthetic systems. Overall, these findings highlight a striking structural variability among GPCR-arrestin signaling complexes.

## Introduction

The biological role of arrestins in G protein-coupled receptor (GPCR) regulation was first discovered in the visual system more than 40 years ago, when arrestin-1 was shown to bind to the light-activated rhodopsin, resulting in the inhibition of receptor signaling (*1, 2*). The first non-visual arrestin, discovered and characterized as a regulator of the β2-adrenergic receptor (β2AR) function (*3*), was named β-arrestin and then β-arrestin1 (βarr1, or arrestin-2). Since then, a wealth of studies have defined the functions of arrestins in regulating GPCR desensitization, endocytosis, and intracellular trafficking (*4, 5*). Beyond these roles, arrestins have been involved in the control of multiple cellular signaling pathways as scaffolding proteins (*6*). One of their better-understood function is to activate mitogen-activated protein kinases (MAPKs), associated with cell cycle regulation, cell growth, and differentiation (*7*). Although it is currently accepted that MAP kinase activation requires endocytosis of stable GPCR-βarr complexes, it was recently shown that βarrs can drive MAP kinase signaling from clathrin-coated structures (CCSs) after GPCR dissociation (*8*). This ‘at a distance’ βarr activation in which transient engagement of the GPCR acts catalytically, also requires a series of interactions with membrane phosphoinositides and CCS-lattice proteins (*9*). The molecular mechanisms and the cellular function of GPCR-βarr interactions were particularly well studied using the arginine-vasopressin (AVP) V2 receptor (V2R) which is involved in the control of water reabsorption and urine concentration in the kidney (*10, 11*). This archetypal model system is well suited to analyze GPCR-βarr assembly due to a long-lasting and stable interaction. Indeed, the V2R binds both βarr1 and βarr2 (or arrestin-3) with similar high-affinity (*12*). Moreover, βarrs remain associated with desensitized V2R during clathrin-mediated endocytosis, a phenomenon directly linked to specific clusters of phosphorylated residues in the receptor C-terminal tail (V2RCter). This sustained interaction was first shown to dictate the slow trafficking of the V2R, in particular its rate of dephosphorylation, recycling, resensitization, and/or degradation (*13, 14*). More recently, this sustained interaction was proposed to enhance AVP-induced cyclic adenosine monophosphate (cAMP) signaling from internalized V2R within endosomes (*15*). The V2R system was also used to investigate how GPCR phosphorylation patterns can orchestrate arrestin conformations and arrestin-dependent distinct signaling pathways (*16*). A synthetic phosphorylated Cter peptide of the V2R (V2Rpp) was shown to functionally and conformationally activate βarr1 (*17*), and its strong affinity was used to investigate the active-state structure of βarr1 using X-ray crystallography (*18*). The complex was captured in the presence of a synthetic antibody fragment Fab30, and revealed at high resolution how the N-lobe of βarr1 accommodates the V2R peptide. βarr1 and V2Rpp make extensive contacts, primarily through charge complementarity interactions between phosphate moieties of V2Rpp and arginine/lysine residues of βarr1 (*19*). The high affinity of V2Rpp for βarr1 was also instrumental in determining the three-dimensional (3D) structures of both muscarinic M2 receptor (M2R)-βarr1 and β1-adrenergic receptor (β1AR)-βarr1 complexes, as in both cases, the natural C-termini of receptors were replaced with that of the V2R (*20, 21*). Both complexes show a V2RCter location similar to that determined in the V2Rpp-βarr1-Fab30 complex.

Although the presence of a phosphorylated V2RCter was necessary for the structure determination of active βarr1 and several GPCR-βarr1 complexes, the structure of the native full-length V2 receptor in complex with βarrs has not been reported yet. Here, we describe the cryo-electron microscopy (cryo-EM) structure of the AVP-bound wild-type human V2R in complex with a truncated form of human βarr1 stabilized by the single-chain variable fragment of Fab30 (ScFv30). Together with the recent structures of the active conformation of the AVP-bound V2R in complex with the Gs protein (*22–24*), these new findings provide major molecular and structural information to better understand arrestin-GPCR interactions as well as V2R-associated signaling pathways.

## Results

### Cryo-EM structure determination of the complex

The AVP-V2R-βarr1ΔCT-ScFv30 complex and the cryo-EM grids were prepared as described in the Materials and Methods section (**Suppl Fig. S1-S3**). A dataset of 14,080 movies was recorded on a Titan Krios microscope for single-particle analysis. To avoid missing any particle, a large number of objects was picked up using two algorithms that take advantage of neural networks and training strategies, and processed with Relion (see Materials and Methods and **Suppl Fig. S4**). Iterative rounds of 2D classification revealed particle classes with clear secondary structural details like V2R transmembrane (TM) domains (**Suppl Fig. S5**). To tackle the strong heterogeneity of the protein complex, a subset of particles sorted from best 2D class averages was imported into CryoSPARC (*25, 26*) and subjected to iterative cycles of two-models ab-initio refinement. The stack of particles selected at each round corresponds to the best-resolved model and also the largest particle stack. The successive models presented the same overall structural organization of the AVP-V2R-βarr1ΔCT-ScFv30 complex (**Suppl Fig. S6**) in which the βarr1ΔCT-ScFv30 complex binds to the detergent micelle and adopts the same relative orientation with respect to V2R (**Suppl Fig. S4, S6**). In the end, this process resulted in a subset of 27,637 particles that once subjected to a nonuniform (NU) 3D refinement, allowed us to compute a density map with global resolution (Fourier shell correlation (FSC) = 0.143) of 4.75 Å for the AVP-V2R-βarr1ΔCT-ScFv30 complex and a 4.2 Å global resolution density map (EMD-14223) for the V2RCter-βarr1ΔCT-ScFv30 after V2R and detergent micelle signal subtraction (**Suppl Fig. S7**). The particles were then further curated resulting in a subset of 8,296 particles which was subjected to a NU 3D refinement, yielding a Cryo-EM map with a global resolution of 4.7 Å (EMD-14221) for the AVP-V2R-βarr1ΔCT-ScFv30 complex (**Suppl Fig. S7**). Data collection and processing are summarized in **Suppl. Table S1**.

The moderate resolution of reconstructions, therefore, indicates a high dynamical behavior of the AVP-V2R-βarr1ΔCT-ScFv30 complex around a preferential structural organization. However, the final EM-maps that display local resolution ranging from 3.5Å to 5.5Å for the AVP-V2R-βarr1ΔCT-ScFv30 (**Suppl Fig. S7C**) and the V2RCter-βarr1ΔCT-ScFv30 complexes (**Suppl Fig. S7D**) allowed us to clearly determine the position and orientation of V2R, βarr1ΔCT, and ScFv30, and to model their backbones (**Fig. 1**). The N-terminus (residues 1-31), parts of the intracellular loops (ICLs) (residues 148-156 from ICL1; residues 183-188 in ICL2; residues 239-263 in ICL3) and the Cter of V2R (residues 343 to 355 and residues 369-371) are not included in the final model. The model of βarr1ΔCT includes residues 6 to 365 whereas ScFv30 is nearly complete (residues 110-128 are missing). AVP and a phosphatidylinositol-4,5-bisphosphate (PtdIns(4,5)P_2_) analogue, the dioctyl-PtdIns(4,5)P_2_ (diC8PIP2), were also constructed in the refined model (**Fig. 1**). Refinement and validation statistics of the AVP-V2R-βarr1ΔCT-ScFv30 and V2RCter-βarr1ΔCT-ScFv30 structural models (PDB 7r0c and 7r0j, respectively) are summarized in **Suppl Table S1**. The overall architecture of the AVP-V2R-βarr1ΔCT-ScFv30 complex (**Fig. 1**) is similar to the one previously described for muscarinic M2 receptor (M2R)-, neurotensin receptor 1 (NTSR1)-and β1-adrenoceptor (β1AR)-βarr1 complexes (*20, 21, 27, 28*). Indeed, the V2R seven TM helical bundle engages the βarr1ΔCT in a core conformation (as defined in (*29*)) with its Cter contacting the βarr1ΔCT N-domain and the ScFv30 (**Fig. 1**). The βarr1ΔCT interacts with V2R through its central crest region whereas its hydrophobic C-edge loop (residues 330-340) inserts into the detergent micelle. The ScFv30 interacts with both βarr1ΔCT N-and C-lobes. The model revealed that both V2R and βarr1ΔCT present characteristics of active state structures with specific features as discussed below.

**Fig. 1.**
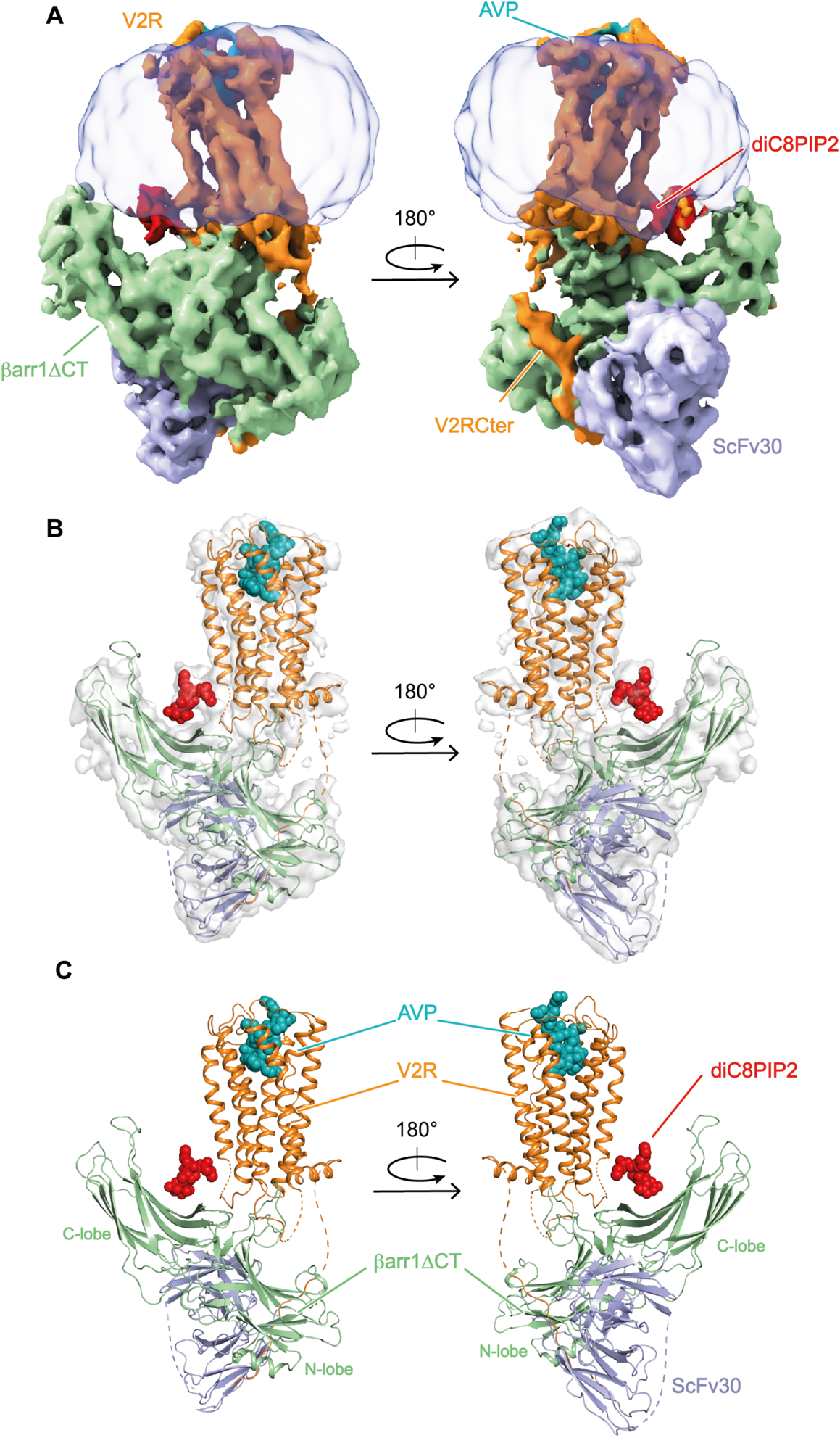
Overall architecture of the AVP-V2R-βarr1ΔCT-ScFv30 complex. (**A**) Orthogonal views of the cryo-EM density map of the complex. V2R and V2RCter are colored in orange, AVP in cyan, βarr1ΔCT in light green, ScFv30 in grey-blue, LMNG detergent micelle in transparent grey. The diC8PIP2 is shown in red. (**B**) Superimposition of the density map and the corresponding model of the complex. Missing parts in the protein chains are shown as dashed lines. The color scheme is as in panel A. (**C**). Final 3D model of the complex.

### The βarr1 engages the V2R with an atypical orientation and a tilted conformation stabilized by the βarr1 C-edge and diC8PIP2

From reported structures of GPCR-βarr1 complexes (*20, 21, 27, 28*), βarr1 can harbor two main perpendicular orientations relative to the GPCR bundle axis, separated by a rotation of approximately 90° parallel to the membrane plane. One is observed for M2R and β1AR complexes and the other for NTSR1 complexes (**Fig. 2A**). Surprisingly, the βarr1ΔCT presents an atypical intermediate position in the AVP-V2R-βarr1ΔCT-ScFv30 complex, being rotated by 54° or 38° when compared to the NTSR1-βarr1 or to the β1AR-βarr1 complex structures, respectively (**Fig. 2A**). A comparison with the other reported GPCR-βarr1 complexes further highlights the unique organization of AVP-V2R-βarr1ΔCT-ScFv30 complex (**Suppl Fig. S8**). Besides the atypical orientation, the C-edge of the βarr1ΔCT C-lobe inserts into the detergent micelle (**Fig. 1A**). This interaction causes a strong tilt with a 55° angle between the longitudinal axis of βarr1ΔCT and V2R, respectively (**Fig. 2B**). Interestingly, βarr1 C-edge membrane anchoring has been observed in the structure of all known GPCR-βarr1 complexes (**Fig. 2B**), whatever the artificial membrane-like environment used for their purification and stabilization (detergent micelles of different chemical nature, nanodiscs). This phenomenon participates in the asymmetry of the different complexes and probably helps in stabilizing βarr1 interactions with GPCRs (*20*). A tilt comparable to the one determined in the AVP-V2R-βarr1ΔCT-ScFv30 structure has been observed in the cryo-EM structure of the NTS_8-13_-NTSR1-βarr1ΔCT complex (*28*), prepared with equivalent detergent micelles (a mix of LMNG, GDN and CHS), and also in the presence of the diC8PIP2 (**Fig. 2B**). In other complexes, in which diC8PIP2 is not present and where the lipid environment of the GPCRs is different (nanodiscs), the arrestin tilt is less pronounced, with an angle ranging from 75 to 60° (M2R-βarr1, β1AR-βarr1 and NTSR1-βarr1 complexes) (**Fig. 2B**). We hypothesize that the βarr1ΔCT-V2R tilt may be amplified due to the small size of these LMNG-GDN-CHS micelles compared to a planar bilayer like the plasma membrane or even nanodiscs (75 versus 55°). As previously described for the NTSR1-βarr1ΔCT complex (*28*), this tilted orientation may reflect the importance of GPCR-arrestin interactions in the context of subcellular structures with a high degree of curvature like CCSs or endosomes (*8, 9*). It is worth noting that although the AVP-V2R-βarr1ΔCT-ScFv30 and the NTS_8-13-NTSR1_-βarr1ΔCT complexes were prepared in the same detergent micelles and in the presence of diC8PIP2, the orientation of βarr1ΔCT relative to the GPCR bundle in these two assemblies is quite different (**Fig. 2A-B and Suppl Fig S8**). This implies that the βarr1ΔCT original orientation in the AVP-V2R-βarr1ΔCT-ScFv30 complex is mainly driven by a peculiar arrestin-GPCR interface, as discussed later in the manuscript.

**Fig. 2.**
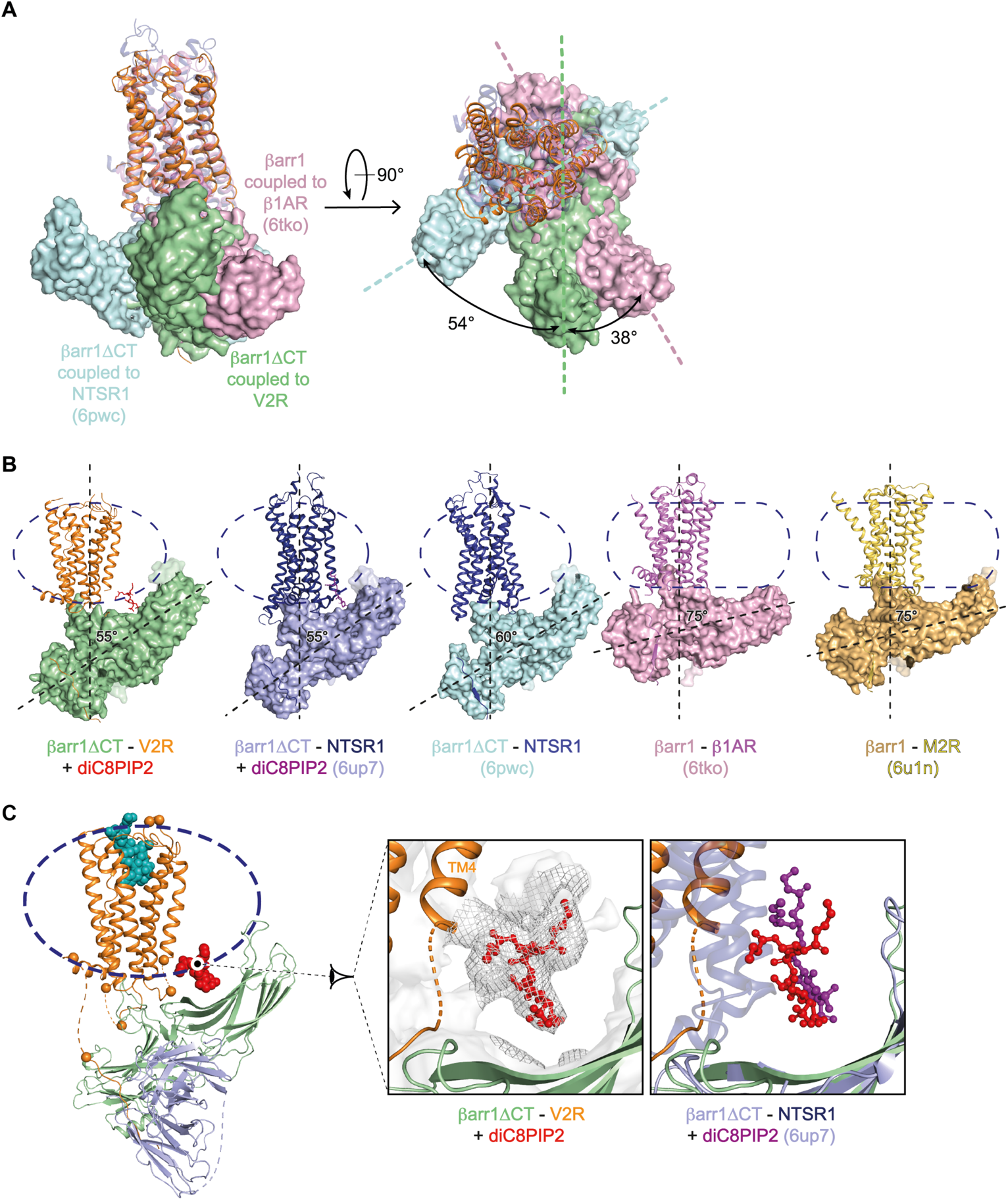
Atypical orientation of βarr1ΔCT and tilted conformation involving the C-edge and diC8PIP2. (**A**) Overlay of V2R-βarr1ΔCT structure with NTSR1-βarr1ΔCT (6wpc) and β1AR-βarr1 (6tko) structures, on the basis of alignment of the receptor chains, viewed from the membrane (left) and from the extracellular space (right). The difference in the orientation of βarr1ΔCT in the complexes is given by the angle of rotation. V2R and βarr1ΔCT are colored like in Fig. 1, βarr1 and β1AR (6tko complex) are in pink, βarr1ΔCT and NTSR1 (6pwc complex) are in blue and purple, respectively. (**B**) Comparison of the tilted conformations of βarr1 (or βarr1ΔCT) in the different GPCR complexes. The contacts between the βarr1 C-edge and the membrane-like environment is visible for each complex. The detergent micelles (V2R, NTSR1 complexes) or nanodiscs (β1AR, M2R complexes) are shown as dashed lines. The angle between the longitudinal axis of βarr1s and GPCRs, respectively, is indicated for each complex. The diC8PIP2 in V2R and NTSR1 (6up7) complexes is shown in red and mauve, respectively. V2R and βarr1ΔCT are colored like in Fig. 1. NTSR1 in both complexes (6up7 and 6pwc) is in dark purple whereas βarr1ΔCT is in purple and blue respectively. βarr1 and β1AR (6tko complex) are in pink, βarr1 and M2R (6u1n complex) are in gold and yellow, respectively. (**C**) 3D model of the AVP-V2R-βarr1ΔCT-ScFv30 complex (left panel). A close-up view of the diC8PIP2 position is shown in the central panel. Its density is displayed as a mesh. Overlay of V2R-βarr1ΔCT with NTSR1-βarr1ΔCT (6up7) (right panel) shows that diC8PIP2 superimposes in the two complexes (alignment is done onto βarr1ΔCT moieties). The color scheme is as in panel B.

A phosphoinositide/phosphoinositol binding site was identified in the C-lobe of βarr1/2 proteins (30) and was shown to play a key role in anchoring these GPCR signaling partners in the plasma membrane at CCSs. In βarr1/2, basic residues, likely to interact with negatively charged phosphates of phosphoinositides (K232, R236, and K250 *versus* K233, R237, and K251 in βarr1 and βarr2, respectively), participate in this binding site. Their mutation leads to βarr1/2 being unable to be recruited neither to CCSs (*30*), nor to the plasma membrane (*9*). Interestingly, a βarr1 variant containing these 3 mutations showed a 40% reduction in recruitment to NTSR1 when compared to the wild-type (*28*). In the NTSR1-βarr1ΔCT complex, diC8PIP2 forms a bridge between the membrane side of NTSR1 TM 1 and 4 and the C-lobe of βarr1ΔCT (**Fig. 2B**).

Our density map (**Fig. 1A-B**) revealed an elongated density protruding from the detergent micelle in a position contacting the βarr1 phosphoinositide binding site, close to residues K232, R236, K250, K324 and K326. We thus modeled a diC8PIP2 molecule in this density (**Fig. 1A-B and Fig. 2C**). Although the orientation of the phosphatidylinositol head group is ambiguous at the resolution of the map, distances between the phosphates 4 and 5 of the inositol ring and these positively charged residues of the C-lobe surface may be compatible with a direct interaction (less than 4 Å if we consider R236 and K324). Moreover, when aligning the AVP-V2R-βarr1ΔCT-ScFv30 and NTS_8-13_-NTSR1-βarr1ΔCT (*28*) complexes onto the βarr1ΔCT protein, the diC8PIP2 moieties also superimpose (**Fig. 2C, right panel**). Due to the atypical orientation of −βarr1ΔCT relative to V2R, the phosphoinositide molecule bridges the C-lobe of βarr1ΔCT with the membrane side of V2R TM4 only (**Fig. 2C, central panel**). The presence of the phosphoinositide moiety may help the insertion of βarr1ΔCT in the detergent micelle and the stabilization of the V2R-βarr1ΔCT interactions, resulting in a tilt similar to that in NTS_8-13_­NTSR1-βarr1ΔCT structure. To analyze this potential stabilizing effect, we performed MD simulations. We found that in the presence of diC8PIP2, root-mean-square deviations and fluctuations of βarr1ΔCT Cα carbons were obviously reduced (**Suppl Fig. S9**). Moreover, the diC8PIP2 constrained βarr1ΔCT C-lobe in a tilted conformation (**Suppl Fig. S10, top**). In addition, βarr1ΔCT exhibited a mobility (rotation and tilting) which was less pronounced in the presence of diC8PIP2 (**Suppl Fig. S9–10**). In conclusion, the presence of diC8PIP2 in the sample combined with insertion of the βarr1ΔCT C-edge into the detergent micelle are probably key parameters to enhance the stability of the AVP-V2R-βarr1ΔCT-ScFv30 complex.

### V2R and βarr1ΔCT display active conformational states

Both the receptor and βarr1 displayed the main hallmarks of active conformations (**Fig. 1 and Fig. 3**). From the receptor point of view, a clear density is observed for the full agonist AVP at the top of the 7TM helix bundle adopting, at this level of resolution, a conformation close to those reported for the active AVP-V2R-Gs structures (**Fig. 3A**) (*22–24*). Both V2R TM6 and TM7 exhibited a displacement by 10 Å and 5 Å respectively, when compared to their counterparts in the inactive antagonist-bound OTR structure (**Fig. 3B**) (*31*). These displacements are similar to those observed in the different structures of AVP-V2R-Gs complexes (*22–24*). From the βarr1 point of view, the superimposition of βarr1ΔCT with the inactive βarr1 clearly shows a rotation of the C-lobe of ∼20 degrees relative to the N-lobe (**Fig. 3C**), as previously described (*32*). Moreover, the alignment of βarr1ΔCT to the active βarr1 demonstrates that the finger loop (FL), gate loop (GL), middle loop (ML), and lariat loop (LL) also adopt an active state to shape a central crest necessary for coupling to GPCRs (**Fig. 3C**). Based on the density map with a better global resolution of 4.2 Å, the different loops of the active form of βarr1ΔCT were confidently assigned and modeled (**Fig. 3D**). The conformation of FL, ML, LL from N-lobe on one side, and that of C-loop (CL) and βstrand 16 from C-lobe on the other side circumscribe a specific furrow (**Fig. 3D**, red arrow) which takes a major part in the specific orientation between the receptor and the βarr1 (see below).

**Fig. 3.**
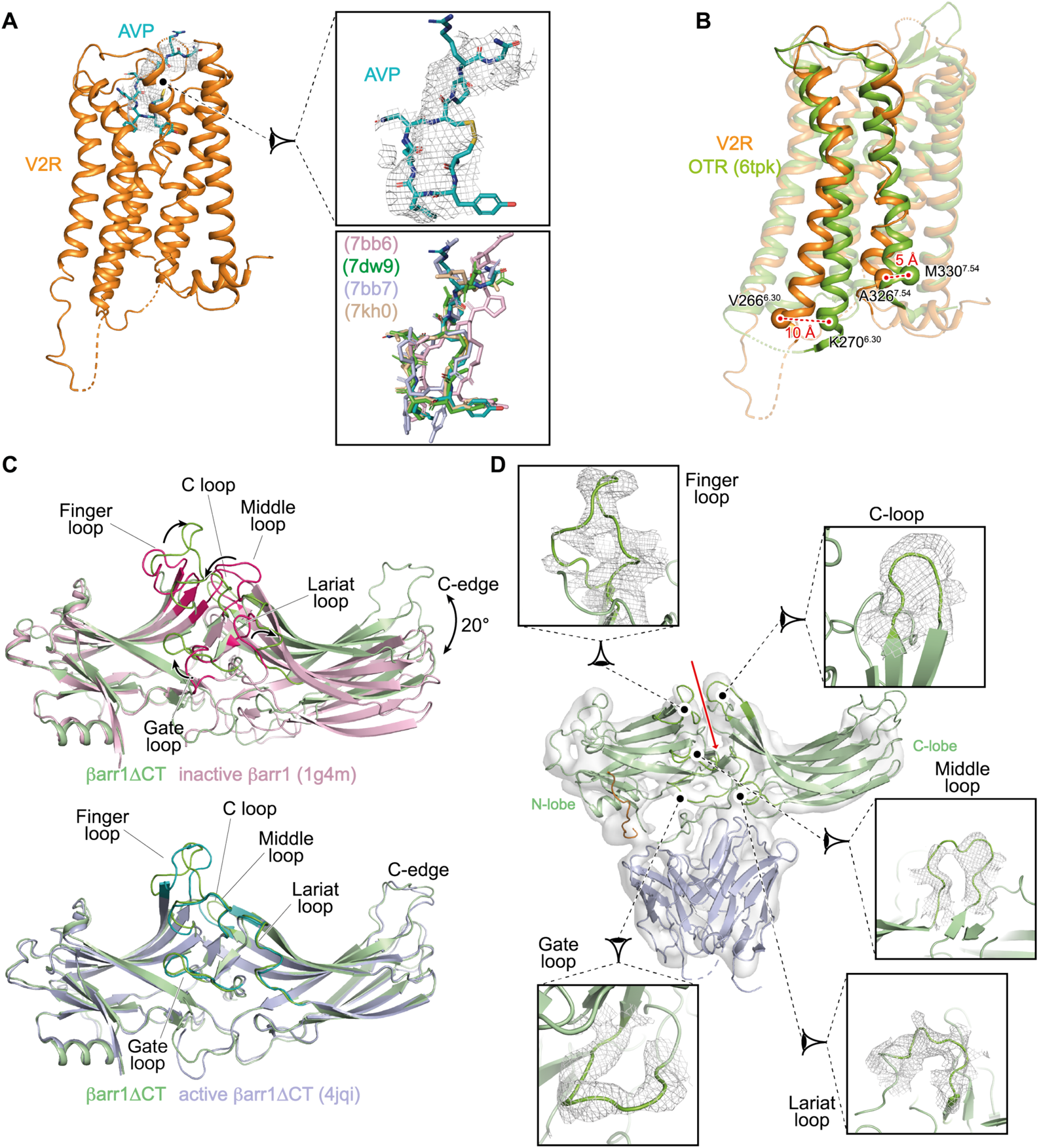
V2R and βarr1ΔCT are in active conformations. (**A**) AVP (blue in the model, density is shown as a mesh) binding to V2R (orange). A close-up view of the AVP binding pose is shown on the right (top panel). Overlapping of V2R-bound AVP in both G protein-and arrestin-associated complexes (bottom panel). (**B**) Comparison of the V2R structure with the inactive OTR (green) structure. Residues 6.30 and 7.54 (Ballesteros-Weinstein numbering) are chosen as references (V266 and A326 in V2R, K270 and M330 in OTR) for measuring the outward (10 Å) and inward (5 Å) movement of TM6 and TM7, respectively. (**C**) The βarr1ΔCT (pale green) in the V2R complex is superimposed onto inactive (top panel) and active (bottom panel) states of the βarr1 (1g4m and 4jqi, respectively). Movements of the different loops (in raspberry in the inactive conformations) are indicated by arrows. The C-lobe is translated by 20° upon activation. Inactive βarr1 is illustrated in pink, active βarr1ΔCT is in light blue. (**D**) Overlay of the density map and the corresponding model of the βarr1ΔCT with close-up views for the different active loops. The color scheme is identical to that of Figs. 1 and 2. Each panel displays the map density as a mesh and the 3D model as a ribbon. The red arrow indicates the central furrow between βarr1ΔCT N-and C-lobes.

### The βarr1/V2R interaction surface defines a novel orientation for an arrestin-GPCR signaling complex

The peculiar architecture of the AVP-V2R-βarr1ΔCT-ScFv30 complex results in an original V2R-βarr1 interface, as compared to those reported for other GPCR-βarr1 complexes. Indeed, all ICLs as well as the 7TM cavity of the V2R are in contact with βarr1ΔCT (**Fig. 4**). First, the density map reveals that ICL1 of V2R directly contacts the ML in the central crest of βarr1ΔCT (**Fig. 4A**). We speculate that G69 and H70 residues of the V2R ICL1, which have been shown to interact with the N-ter helix of Gs α subunit (*22*), are probably involved in this interaction. Second, although ICL2 is not entirely seen in the density map, this V2R region (residues R139 to A147) strikingly binds in a particularly well-defined furrow between the N-and C-lobes of βarr1ΔCT, lying in a central position (**Fig. 3D, Fig. 4A-B**). Third, most of the V2R ICL3 is not resolved (residues 239-263 are missing), however clear contacts are seen with the N-lobe of βarr1ΔCT, in particular with the N-terminal part of the FL (**Fig. 4A**). Due to the orientation of βarr1ΔCT relative to V2R, we speculate that V2R ICL3 may transiently interact with a region defined by the FL, the β strands of the N-lobe and the 160-loop (the β strands 9 and 10 junction). In addition, the ICL3 of V2R displays a cluster of arginine residues (RRRGRR at positions 247-252) that may also create ionic contacts with βarr1ΔCT negatively charged residues, for instance E66, E67, D78, D143, E134, E145, E152, E155, E156. To explore these potential interactions, we have performed molecular dynamics (MD) simulations of AVP-V2R-βarr1ΔCT including V2R ICL2 and ICL3. These two ICLs were highly mobile in the different simulations (**Suppl Fig. S10**), consistent with the lack of visibility in the cryo-EM density maps. Despite the high mobility of V2R ICL3 in the simulations, persistent interactions were only observed between the RRRGRR arginine cluster and residues V127-E134 in the ML of βarr1 N-lobe (**Suppl Fig. S9, S10 and Suppl Table 2**).

**Fig. 4.**
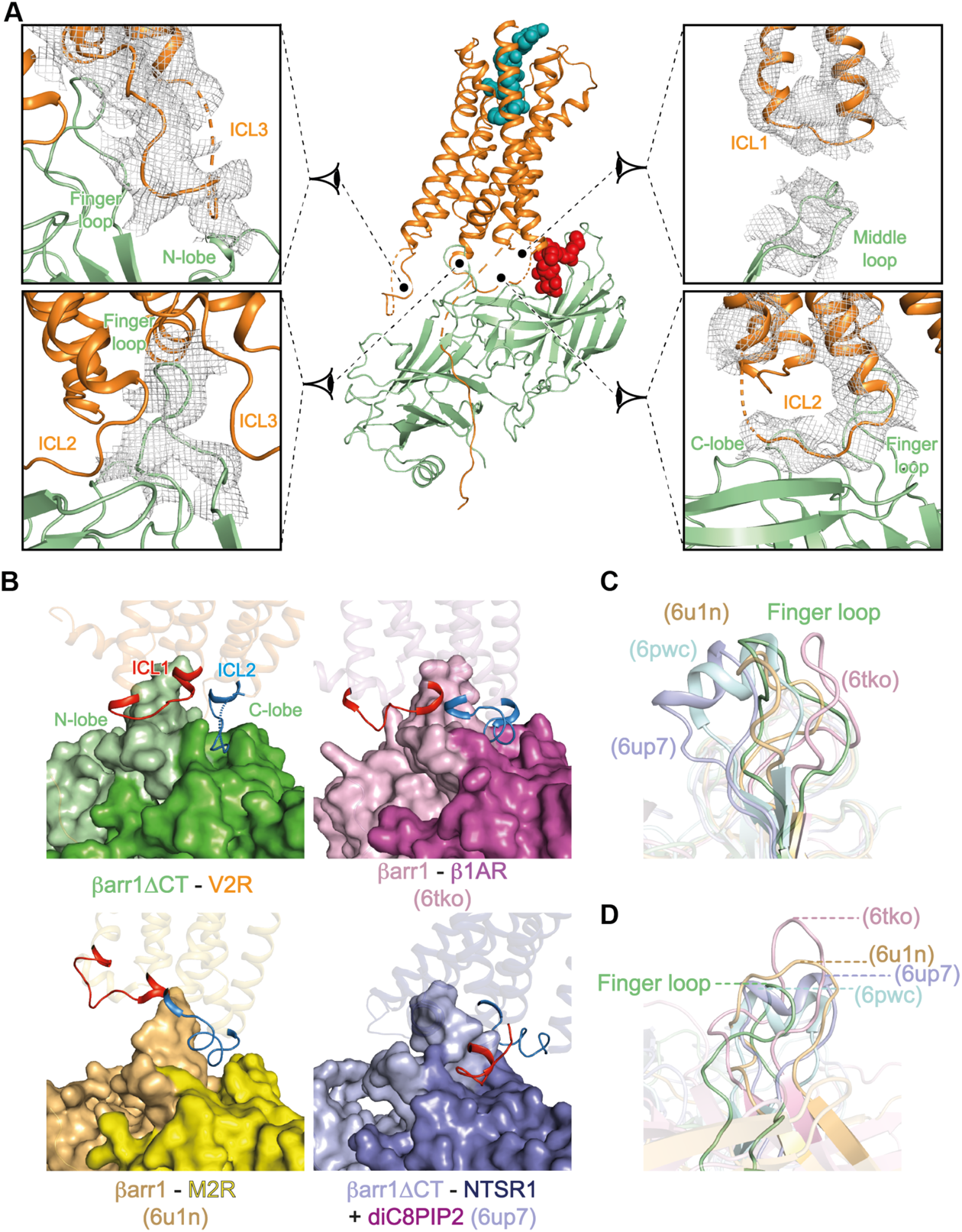
The V2R/βarr1ΔCT interface. (**A**) The model of the AVP-V2R-βarr1ΔCT complex is shown in the middle with close-up views for the different domains involved in the interaction surface. The color scheme is identical to that of Fig. 1. Each zoom panel displays the map density as a mesh and the corresponding model as a ribbon. (**B**) Position of ICL2 (or ICL1) in the central furrow, for V2R-, β1AR-, M2R-and NTSR1 (PDB 6up7)-associated complexes. ICL1 is in red, ICL2 is in blue. To have a better view of the central crevasse, the N-lobe and the C-lobe of βarr1s are highlighted in light and dark related colors: green for the present V2R-βarr1ΔCT complex, pink for 6tko, yellow for 6un1 and purple for 6up7. (**C**) Orthogonal views of the βarr1 FL in the different complexes. The color scheme is identical to that of Fig. 2. The top panel shows alignment made with βarr1s in order to illustrate the FL variable conformations in the different GPCR complexes, whereas the bottom panel shows alignment made with GPCRs to highlight the FL insertion deepness.

Altogether, as compared to the structures of other GPCR-βarr1 complexes, the combination of interactions between ICLs of V2R and the βarr1ΔCT is quite singular. Indeed, as opposed to V2R, ICL1 from M2R does not interact at all with βarr1 (**Fig. 4B**) while the one of β1AR is positioned close to both ML and part of FL (**Fig. 4B**). Interestingly, the ICL2s from β1AR or M2R also insert into the central furrow, but with an orientation and deepness which are slightly different (**Fig. 4B**). Surprisingly for the NTSR1-βarr1 complexes, ICL1 binds to this furrow instead of ICL2 (**Fig. 4B**), a specificity that may explain the particular rotation (∼90°) of βarr1 in the NTSR1-βarr1 structure when compared to the β1AR and M2R complexes (**Fig. 2A**). The structures of β1AR-βarr1 and M2R-βarr1ΔCT complexes (*20, 21*) show that residues in the βstrand 16 (Y249), CL (I241), LL (R285) or FL (Y63, R65) domains of βarr1 directly participate in the binding of ICL2 and consequently may affect the orientation between the receptor and βarr1. Of note, Y249 and R285 also interact with ICL1 in the NTSR1-βarr1 complexes (*27, 28*). Although the global resolution is lower for the V2R-βarr1ΔCT complex, we speculate that most of these residues are involved in the binding of ICL2. Taken together these data suggest that depending on the GPCR, the interactions between ICL1 or ICL2 with the particular central furrow of βarr1 are, at least in part, involved in determining the architecture of the signaling GPCR-βarr1 complexes.

Finally, the V2R transmembrane cavity formed by the outward motion of TM6 engages the FL of βarr1ΔCT (from Y63 to L73) (**Fig. 4C** and **Suppl Fig. S1B**). A comparison of the V2R-coupled βarr1ΔCT structure with those published for other complexes highlights a strong variability of FL, both in its conformation and its GPCR binding deepness (**Fig. 4C**). In all GPCR-βarr1 complexes, the FL inserts the receptor core in a pocket delimited by TM2, 3, 6, 7, and establishes contacts with different residues from these TM helices. However, depending on the receptor, FL can adopt an α-helical domain (in the NTSR1-βarr1ΔCT complex) or can be deeply inserted in the GPCR core (in the β1AR-βarr1 complex), contacting conserved I^6.40^ or Y^7.53^ (Weinstein-Ballesteros nomenclature) (**Fig. 4C**). At the V2R-βarr1ΔCT interface, the FL inserts V2R like in other GPCR-arrestin complexes, but not as deep into the TM core as observed for βarr1 FL in β1AR (*21*). More particularly, it seems to contact residues from the cytoplasmic sides of TM 2, 3, and 6. At this interface, D69, positioned at the tip of FL, could establish an ionic bridge with R137^3.50^ (Ballesteros-Weinstein nomenclature) in V2R TM3. This interaction has also been seen in the M2R-βarr1 and the β1AR-βarr1 complexes (*20, 21*), strengthening this hypothesis. Interestingly, R137^3.50^ is part of the ionic lock motif (broken in the active V2R conformation) and has been shown to directly interact with the free carboxylic acid function of the Gs protein α subunit Cter in the active structure of the AVP-V2R-Gs-Nb35 complex (*22*).

Therefore, in addition to the key ICL2-βarr1ΔCT interactions, additional structural parameters including ICL1 positioning, ICL3 dynamic interaction, and a particular FL conformation and binding deepness in the 7TM core, are involved in defining the atypical orientation of βarr1ΔCT bound to the V2R.

### Constitutive phosphorylation of V2R and interaction with βarr1

The Cter of GPCRs plays a key role in many aspects of their regulation, through phosphorylation by various GPCR kinases (GRKs) and subsequent binding to arrestins. The number and arrangement of phosphates may vary substantially for a given receptor and different phosphorylation patterns have been shown to trigger different arrestin-mediated effects (*33*). The structural basis regulating this phosphorylation barcode has been revealed recently (*16*), indicating how GPCR phosphorylation affects arrestin binding and conformation.

Clear interactions between the Cter of V2R and the N-lobe of βarr1 are visualized in the density map of the locally-refined βarr1ΔCT-ScFv30 subcomponents (**Fig. 5A**). This region of the complex displays better densities than elsewhere with local resolution between 3.5 and 4Å (**Suppl Fig. S7**). Densities for V2R phosphoresidues S357, T359, T360, S362, S363 and S364 could be identified (**Fig. 5A**), in agreement with their position defined in the active structure of βarr1 in complex with a chemically-synthesized V2Rpp (*18*) or with that seen in the M2R-βarr1 and β1AR-βarr1 complexes (both having a V2R Cter fused instead of their natural Cter). To confirm the presence of these phosphoresidues in the V2RCter, phosphoproteomics of the purified V2R has been performed using trypsin cleavage and LC-MS/MS approach allowing to identify phosphopeptides and to determine the phosphosite localization (significant phosphosites are correlated to a localization probability superior to 0.75). Most of the phosphoresidues identified in the density map are indeed phosphorylated (**Suppl Fig. S11**), since S357, T359, S362, and S364 present a localization probability above 0.75. In addition, T347 and S350, positioned right after the helix8 also display a high localization probability (but are not seen in the density map). Interestingly, phosphoproteomics experiments also identified phosphorylation sites in ICL3 (S241, T253 and S255 which are however not visible in the density map) (**Suppl Fig. S11**), but we did not observe apparent interactions between these phosphorylated residues and βarr1ΔCT in MD simulations (**Suppl Table S2**). Surprisingly, a majority of the residues that are phosphorylated (both in the ICL3 and in the Cter), were post-translationally modified whether the V2R-expressing Sf9 cells were stimulated or not with the full agonist AVP (1 µM for 30 min) before harvesting (**Suppl Fig. S11**). This means that the V2R is constitutively phosphorylated in the Sf9 insect cells. The presence of GRKs has been studied in this cellular system and the role of these insect cell GRKs has been proven in the agonist-induced desensitization and phosphorylation of the human M2 muscarinic and serotonin 5HT_1A_ receptors (*34, 35*).

**Fig. 5.**
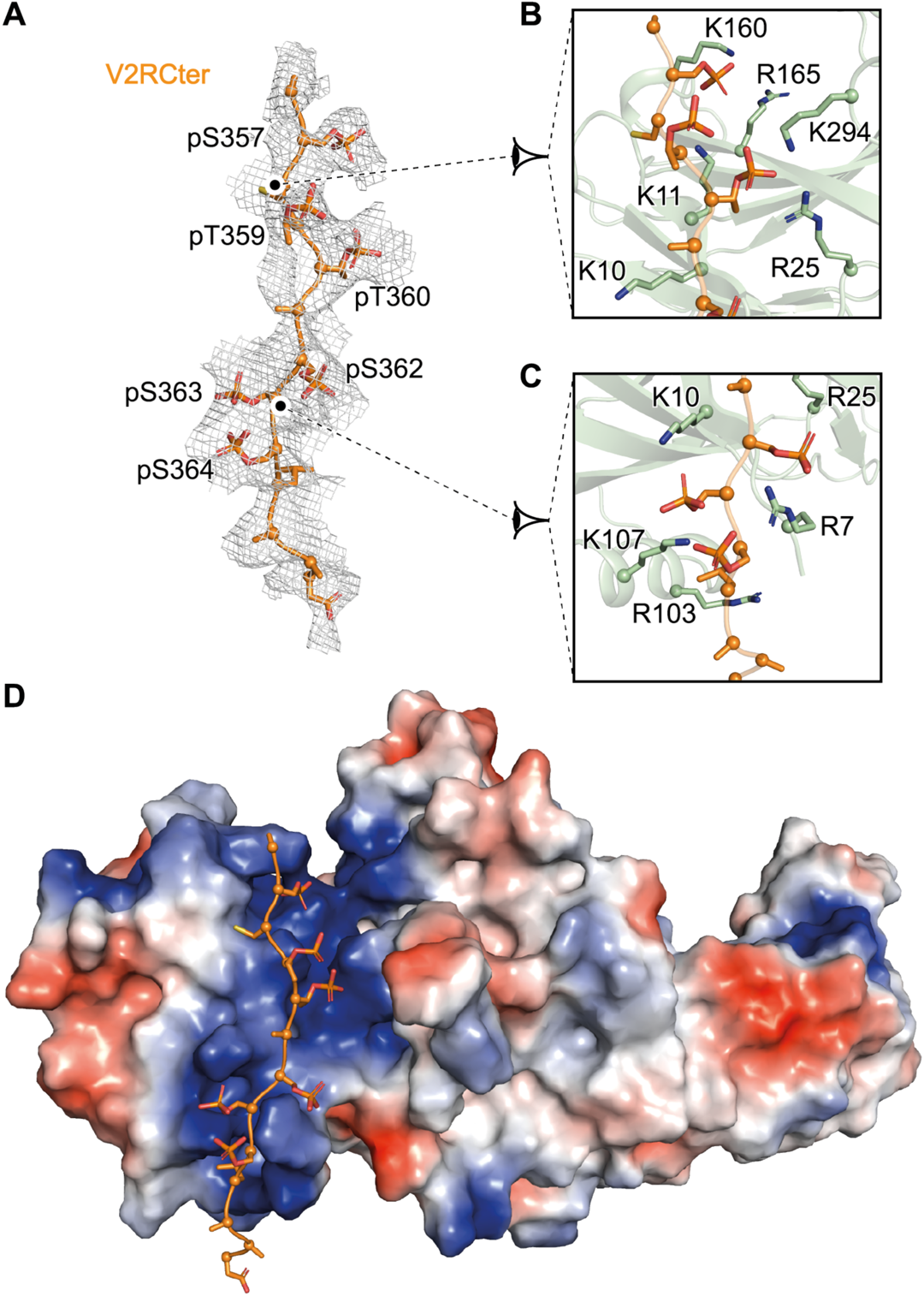
Phosphorylation of V2RCter and interaction with βarr1ΔCT N-lobe. (**A**) Overlay of the cryo-EM map of the V2RCter (mesh) with its corresponding model (in orange). Phosphate moieties of V2R pS357, pS359, pT369, pS362, pS363, and pS364 are shown in red. (**B**) and (**C**) Close-up views of these phosphorylated residues of the V2RCter and of positively charged residues (in pale green) of the N-lobe of βarr1ΔCT. (**D**) Global view of the phosphorylated V2RCter interacting with the N-lobe of βarr1ΔCT. The charge potential surface of βarr1ΔCT is shown in red (negatively-charged residues) and blue (positively-charged residues) representation.

Based on the density map, the phosphoproteomics results and previous structural data, we confidently built the interactions between negatively charged phosphates of the V2RCter residues and positively charged K/R residues of the βarr1 N-lobe (distances less than 4 Å) (**Fig. 5B, C and D**). Phosphates of V2R S357, T359 and T360 are in close proximity to K11 and R25, whereas phosphates of V2R S362, S363 and S364 may establish ionic contacts with R7, K10 and K107 of βarr1 (**Fig. 5B and C**), like in the crystal structure of the βarr1-V2Rpp-Fab30 (*18*), and in the structure of the M2R-βarr1 and β1AR-βarr1 complexes where both GPCR have a V2RCter fused instead of their natural sequence (*20, 21*).

### Comparison of AVP-V2R-Gs-Nb35 and AVP-V2R-βarr1ΔCT-ScFv30 complexes

We recently determined the structure of the AVP-bound V2R-Gs complex (*22*), which led us to compare it with the AVP-bound V2R-βarr1 complex (**Fig. 6A**). Importantly, for both complexes, the interface between V2R and the signaling partners is native and was not modified by protein engineering. Indeed, no mutations were introduced, no chimeric or fusion proteins were constructed, no crosslinking approach was done, no NanoBIT tethering strategy was used (*36*). First, when comparing the structures of AVP-V2R-Gs (PDB 7bb7) and AVP-V2R-βarr1ΔCT complexes to the inactive structure of the related oxytocin receptor (*31*), V2R TM6 moves outward in both cases with a similar range of distance, 13 Å (AVP-V2R-Gs assembly) versus 10 Å (AVP-V2R-βarr1ΔCT complex, **Fig. 3B**), leading to the opening of the receptor core cavity. Second, superposition of V2R from the two complexes (r.m.s.d. 3.5 Å for 1,767 atoms) shows that both Gs and βarr1 insert into this core cavity (**Fig. 6B**). More precisely, the Gs protein α subunit Cter (h5 helix) and the tip of the βarr1 FL overlap in the same space (**Fig. 6B**). Based on the density maps and the corresponding models (**Fig. 6A-B**), the free carboxylic acid function of the Gs protein α subunit Cter (*22*) and the carboxylic function of the D69 of the βarr1 FL may both interact with the R137^3.50^ **(Fig. 6C**), which is part of the ionic lock, as discussed above. The destabilization of this motif is important for the receptor to reach active conformations and to engage with the different G protein-and βarr-dependent signaling pathways. Overall, the V2R conformation in the AVP-bound V2R-Gs complex is very similar to that in the AVP-bound V2R-βarr1ΔCT complex (**Fig. 6B**), but some key domains might present slight differences like the TM7-H8 hinge. These key domains may be involved in biased ligand-induced differential conformations. For instance, using fluorescence spectroscopy and full *versus* biased ligands, the TM7-H8 hinge region was shown to favor βarr1 interaction over G protein (*37*).

**Fig. 6.**
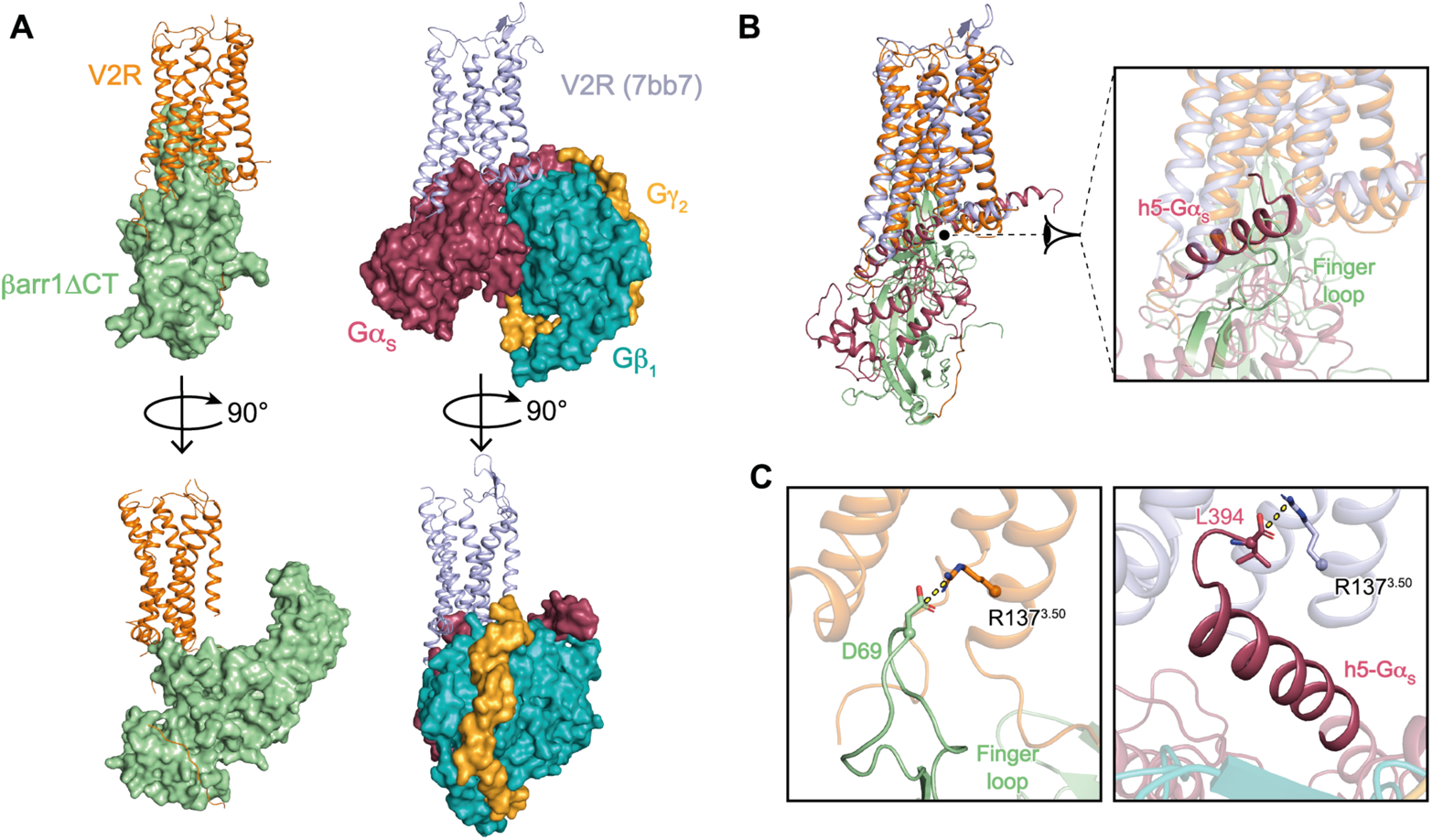
Comparison of AVP-V2R-Gs and AVP-V2R-βarr1ΔCT complexes. (**A**) Orthogonal views of V2R-βarr1ΔCT (left) and V2R-Gs (right, PDB 7bb7) complexes. The color scheme for the V2R and βarr1ΔCT is as in Fig. 1. In the other complex, V2R is shown in blue-grey, Gα_s_ in raspberry, Gβ_1_ in turquoise, and Gγ_2_ in yellow. (**B**) Superimposition of the two complexes with a close-up view at the interfaces. The C-terminal helix of the Gα subunit of the Gs protein (h5-Gαs helix) and the tip of βarr1ΔCT finger loop insert into the transmembrane core cavity of the V2R and overlap in the same binding space. (**C**) In the AVP-V2R-βarr1ΔCT complex, the FL residue D69 carboxylic function may interact with V2R R137 through an ionic bridge. In the AVP-V2R-Gs complex, the ionic interaction between the Cter free carboxylic function of the Gαs with V2R R137 is shown for comparison.

Because each signaling protein is tightly associated with the V2R transmembrane cavity in both assemblies, it is obvious that Gs and βarr1 cannot couple simultaneously when βarr1 is in the “core” conformation (**Fig. 6A**). These observations support a steric hindrance-based desensitization mechanism through competition for an overlapping interface at the cytoplasmic transmembrane surface of the V2R (*38, 39*). This is in agreement with studies showing that desensitization of G protein signaling (i.e. arresting the G protein signaling) is performed exclusively by the receptor core-engaged βarr1 (*29, 40*), whereas a βarr1 in a “tail” conformation is fully capable of performing other canonical functions (i.e., signaling and receptor internalization). Interestingly, simultaneous coupling of the two signaling proteins has also been demonstrated when arrestin is pre-associated in the “tail” conformation, i.e. when the βarr1 only attaches to the phosphorylated Cter of the V2R (*38*). The formation of this megacomplex was proposed to provide a biophysical basis for sustained endosomal G protein signaling (*41*). Its structure has been solved recently (*39*). The AVP-V2R-βarr1ΔCT signaling complex described here only concerns the GPCR core-engaged βarr1, and might be representative of the desensitization step of G protein primary signaling regulated by the βarr1 at the plasma membrane.

## Conclusion

The architecture of the AVP-V2R-βarr1ΔCT-ScFv30 complex described herein notably puts forward an original positioning of the βarr1, an unusual V2R/βarr1 interface and a strong tilt of βarr1ΔCT relative to V2R. This further highlights a striking structural variability among GPCR-arrestin signaling complexes. Although there are multiple sites of interactions between the V2R and βarr1ΔCT, as well as a structurally important interface between the micelle and the βarr1 C-edge, and despite the addition of stabilizing partners such as the ScFv30 and the diC8PIP2, this complex displays a highly flexible behavior. This is in agreement with multiple conformations of arrestins and their versatile role in biological signaling systems (*42*).

The AVP-V2R-βarr1 complex structure is an additional step to better understand receptor conformational changes upon binding to different signaling proteins. In the future, it would be crucial to determine how the structures of V2R-Gs and V2R-βarr1 complexes in the presence of biased ligands are conformationally different than those defined for unbiased AVP. Indeed, developing ligands able to discriminate the Gs protein-and βarr-dependent signaling pathways associated to V2R activation is of crucial importance regarding polycystic kidney disease (*43*) or two V2R-associated genetic diseases with opposite clinical outcomes, congenital nephrogenic diabetes insipidus (*44*) and nephrogenic syndrome of inappropriate antidiuresis (*45*). Structure-based development of novel molecules able to differentiate Gs protein-, “core “ βarr1-or “tail” βarr1-associated V2R conformations will pave the way to design better drugs against kidney pathologies (*46*).

## Acknowledgments

We thank the cryo-EM staff at EMBL of Heidelberg (Germany), the Institut de Génomique Fonctionnelle Arpege Pharmacology (http://www.arpege.cnrs.fr) and Functional Proteomics (http://www.ppm.cnrs.fr) platforms. We thank Perkin Elmer CisBio for providing reagents.

## Author contributions

J.B. and A.F. purified V2R and AVP-V2R-βarr1ΔCT-ScFv30 complexes, screened samples by NS-EM and Cryo-EM, prepared grids, collected and processed Cryo-EM data, generated the cryo-EM maps, and built supplementary figures. H.O. managed the Sf9 cell culture and baculovirus infections, expressed and purified V2R, purified AVP-V2R­βarr1ΔCT-ScFv30 complexes, and prepared grids for Cryo-EM. S.T. constructed the 3D models of the AVP-V2R-βarr1ΔCT-ScFv30 complex. X.C. performed all MD simulations. S.F. produced and purified the βarr1ΔCT. J.S.-P. expressed and purified the ScFv30. J.L.-K.-H. participated in the screening of samples by NS-EM and Cryo-EM. S.U. managed and performed the phosphoproteomic experiments and analysis. N.S. participated in the production and purification of the βarr1ΔCT. R.S. designed the βarr1ΔCT and ScFv30 constructs, and built all principal figures. B.M. designed the V2R construct. S.G., B.M., and P.B. wrote the manuscript with the input from J.B. and A.F. Last, S.G., B.M., and P.B. supervised the project.

## Data and materials availability

The Cryo-EM density maps for the AVP-V2R-βarr1ΔCT-ScFv30 complex have been deposited in the Electron Microscopy Data Bank (EMDB) under accession codes EMD-14221 and EMD-14223. The coordinates for the corresponding models of the AVP-V2R-βarr1ΔCT-ScFv30 complex have been deposited in the Protein Data Bank (PDB) under accession numbers 7r0c and 7r0j. All relevant data are available from the authors and/or included in the manuscript or in the Supplementary Materials.

## Supplementary Materials

Materials and Methods

Figs. S1 to S11

Tables S1 and S2

References (*47-83*)

## Supplementary Materials

### Materials and Methods

#### Data analysis and figure preparation

Figures were created using the PyMOL 2.3.5 Molecular Graphics System (Schrödinger, LLC) and the UCSF Chimera X 0.9 package (*47*). Data were plotted with GraphPad Prism 9.1.1 (GraphPad Prism Software Inc.).

#### V2R expression and purification

The optimized sequence of the human V2R was cloned into the pFastBac1 vector (Thermo Fisher Scientific) using EcoRI/XbaI restrictions sites to enable insect Sf9 cells infection using a baculovirus cell expression system. Since it has been demonstrated that the whole C-terminus of V2R is crucial for arrestin interaction (*13*), it was conserved native in our construct. Indeed, the hemagglutinin signal peptide (MKTIIALSYIFCLVFA), a first Flag-tag (DYKDDDDA), a Twin-Strep-tag (WSHPQFEKGGGSGGGSGGGSWSHPQFEK), a human rhinovirus 3C (HRV3C) protease cleavage site, and a second Flag-tag were all inserted in-frame in the V2R N-terminus to facilitate expression and purification of the receptor (**Suppl Fig. S1A**). N22 was substituted with a glutamine residue to avoid *N*-glycosylation. M1 and L2 residues of the wild-type V2R sequence were not present in this construct. Before production in Sf9 insect cells, this construct was first validated in human embryonic kidney cells to control that it retained wild-type pharmacological and functional properties. First, Kd of the fluorescently-labelled antagonist was 4.22 ± 0.7 nM (n=3) in agreement with that defined for the wild-type V2R (*48*). Then, AVP binding, accumulation of cytosolic cAMP and recruitment of βarrestin2 assays (see below for description of the three methods) all confirmed the V2R is fully functional (**Suppl Fig. S2**). Indeed, the Ki for AVP was 3.17 ± 0.97 nM (n=3), and the Kact for AVP-induced cAMP production and arrestin recruitment were 0.22 ± 0.09 nM (n=3) and 2.02 ± 0.28 nM (n=4) respectively, in accordance with those determined for a wild-type V2R (*22, 48, 49*).

The V2R was expressed in Sf9 insect cells using the Bac-to-Bac baculovirus expression system (Thermo Fisher Scientific) according to the manufacturer’s instructions as previously described (*22*). Briefly, insect cells were grown in suspension in EX-CELL 420 medium (Sigma-Aldrich) to a density of 4 × 10^6^ cells/ml and infected with the recombinant baculovirus at a multiplicity of infection of 2 to 3. The culture medium was supplemented with the V2R pharmacochaperone antagonist tolvaptan (TVP) (Sigma-Aldrich) at 1 µM to increase the receptor expression levels (*50, 51*). The cells were infected for 48 to 54 hours at 28 °C, and expression of the V2R was checked by immunofluorescence using an anti-Flag M1 antibody coupled to Alexa Fluor 488. In order to favor V2R phosphorylation, cells were treated with 1 µM AVP 30 min before being harvested by centrifugation (two steps for 20 min at 3,000 *g*), and pellets were stored at −80 °C until use.

The first step of V2R purification was performed as already described (*22*). Briefly, the cell pellets were thawed and lysed by osmotic shock in 10 mM tris-HCl (pH 8), 1 mM EDTA buffer containing iodoacetamide (2 mg/ml, Sigma-Aldrich), 1 µM TVP, and protease inhibitors [leupeptine (5 µg/ml) (Euromedex), benzamidine (10 µg/ml) (Sigma-Aldrich), and phenylmethylsulfonyl fluoride (PMSF) (10 µg/ml) (Euromedex)]. After centrifugation (15 min at 38,400 *g*), the pellet containing crude membranes was solubilized using a glass dounce tissue grinder (15 and 20 strokes using A and B pestles, respectively) in a solubilization buffer containing 20 mM tris-HCl (pH 8), 500 mM NaCl, 0.5 % (w/v) *n*-dodecyl-β-d-maltopyranoside (DDM, Anatrace), 0.2 % (w/v) sodium cholate (Sigma-Aldrich), 0.03 % (w/v) cholesteryl hemisuccinate (CHS, Sigma-Aldrich), 20 % glycerol, iodoacetamide (2 mg/ml), biotin BioLock (0.75 ml/liter, IBA), 1 µM TVP, and protease inhibitors. The extraction mixture was stirred for 1 hour at 4 °C and centrifuged (20 min at 38,400 *g*). The cleared supernatant was poured onto an equilibrated Strep-Tactin resin (IBA) for a first affinity purification step. After 2 hours of incubation at 4 °C under stirring, the resin was washed three times with 10 column volume (CV) of a buffer containing 20 mM tris-HCl (pH 8), 500 mM NaCl, 0.1 % (w/v) DDM, 0.02 % (w/v) sodium cholate, 0.03 % (w/v) CHS, and 1 µM TVP. The bound receptor was eluted in the same buffer supplemented with 2.5 mM desthiobiotin (IBA). The HRV3C protease was added for overnight cleavage at 4 °C (a 1:20 (HRV3C:V2R) weight ratio). After digestion, the eluate was loaded onto a M2 anti-Flag affinity resin (Sigma-Aldrich). After loading, the DDM detergent was then gradually exchanged with Lauryl Maltose Neopentyl Glycol (LMNG, Anatrace) and glyco-diosgenin (GDN, Anatrace). The LMNG concentration was then decreased gradually from 0.5 to 0.02 %, and that of GDN from 0.125% to 0.005%. The V2R was eluted in 20 mM HEPES (pH 7.5), 100 mM NaCl, 0.02 % LMNG, 0.005% GDN, 0.002 % CHS, 10 µM AVP (Bachem), and Flag peptide (0.4 mg/ml, Covalab). After concentration using a 50-kDa molecular weight cutoff (MWCO) concentrator (Millipore), the V2R was purified by SEC using a Superdex 200 increase (10/300 GL column) connected to an ÄKTA purifier system (GE Healthcare). Fractions corresponding to the pure monomeric receptor were pooled (∼2 ml) and concentrated to 50 to 100 µM with an excess of AVP (200 µM).

#### β-arrestin1 expression and purification

A truncated version of β-arrestin1 at residue 382 (βarr1ΔCT) was produced and purified (**Suppl Fig. S1B**), since it has been shown to display a constitutive activity in cells (*52*). Indeed, this βarr1 variant was able to effectively desensitize β2AR and δ opioid receptor in Xenopus oocytes. In addition, its recombinantly purified version was demonstrated to stably interact with the purified NTSR1 (*28*). It was prepared as follows. BL21(DE3) competent E. coli cells (ThermoFisher Scientific) were transformed using a pET plasmid containing an optimized version of β-arr1ΔCT fused to a Twin-Strep-tag sequence at its N-terminus (NcoI/XhoI subcloning), and large-scale cultures were grown in LB + kanamycin at 37 °C (170 rpm) until an optical density (OD600) at 0.6 U was reached. Cells were induced at 37 °C for 5 hours by adding 0.025 mM IPTG. Cells were collected by centrifugation (two steps for 20 min at 3,000 g), and pellets were stored at −80 °C until use. Cells were resuspended in lysis buffer (20 mM Tris-HCl (pH 8), 1 mM EDTA, 200 mM NaCl, 1 mM β-mercaptoethanol) supplemented with protease inhibitors [leupeptine (5 µg/ml), benzamidine (10 µg/ml), and PMSF (10 µg/ml)]. Cells were lysed by sonication and the lysate was supplemented with MgCl_2_ (5 mM final) and Benzonase (2000 Units). After centrifugation (20 min, 4 °C, 38,400 g), the supernatant was supplemented with biotin BioLock (0.75 ml/liter) and loaded to Strep-Tactin affinity resin at 4 °C. The resin was washed with 20 CV of wash buffer (20 mM Tris pH 8, 200 mM NaCl, 100 µM TCEP). The protein was then eluted with 5 CV of wash buffer supplemented with 2.5 mM desthiobiotin (IBA). Subsequently, it was subjected to a Superdex 200 increase gel filtration step (10/300 GL column) with a buffer containing 20 mM HEPES (pH 7.5), 200 mM NaCl and 100 µM TCEP. The fractions corresponding to the purified βarr1ΔCT were collected, concentrated to approximately 250 µM using a 10-kDa MWCO concentrator (Millipore). Aliquots were then flash-frozen and stored at −80 °C until use.

#### ScFv30 expression and purification

The single chain variable fragment ScFv30 with a Twin-Strep-tag added at its C-terminus was used in this study to enable a stable interaction between the receptor and βarr1ΔCT (*18, 53*). Moreover, this antibody fragment has been shown to not bind to the C-terminus of βarr1 and the corresponding Fab30 was shown to lock the βarr1 in its active conformation (*21*). Briefly, the optimized nucleotide sequence of ScFv30 was cloned in NcoI/PmeI restriction sites of a modified pMT/BIP/V5 vector (Life Technologies) in which the V5 epitope and the 6-His tag were replaced by an enterokinase cleavage site followed by a Twin-Strep-tag (WSHPQFEKGGGSGGGSGGGSWSHPQFEK). This vector is adapted to secreted expression of recombinant proteins in *Drosophila melanogaster* S2 Schneider cell cultures (*54, 55*). In this plasmid, the ScFv30 sequence is in frame with the BIP signal sequence and the Twin-Strep-tag. The expression and purification of ScFv30 were performed as follows. S2 Schneider cells (Life technologies), cultured in serum-free insect Xpress medium (Lonza), were transfected as reported previously (*56*), amplified, and ScFv30 expression was induced with 4 µM CdCl_2_ at a density of ∼10 x 10^6^ cells per ml for 6-8 days for large-scale production.

Cells were harvested by centrifugation (38,400 g, 10 min, 4 °C) to remove cells and cellular debris. Then, protease inhibitors [leupeptine (5 µg/ml), benzamidine (10 µg/ml), and phenylmethylsulfonyl fluoride (PMSF) (10 µg/ml)] were added to the supernatant. The sample was filtered and concentrated at 4 °C using a Vivaflow 200 cassette with a 10-kDa MWCO (Sartorius). When the volume was reaching around 100 ml, Tris-HCl (pH 8, 100 mM) and 0.5 ml Biolock (IBA) were added. The ScFv30 was purified by Strep-Tactin affinity chromatography (IBA) in a buffer containing 100 mM tris-HCl (pH8), 150 mM NaCl, 1 mM EDTA. The eluate was concentrated to reach a 5 to 10 ml volume and dialyzed in 2 steps (O/N at 4 °C, then 2 hours) in a buffer containing 20 mM HEPES pH7.5, 100 mM NaCl. The dialyzed ScFv30 was concentrated to approximately 200 µM using a 10-kDa MWCO concentrator (Millipore). Aliquots were flash-frozen in liquid nitrogen and stored at −80 °C until use.

#### Purification of the AVP-V2R-βarr1ΔCT-ScFv30 complex

As indicated above, all components of the complex (V2R, βarr1ΔCT and ScFv30) were first expressed and purified separately. Then they were mixed in the presence of an excess of AVP (**Suppl Fig. S3**). The mixture was complemented with a phosphatidylinositol-4,5-bisphosphate analogue, the dioctyl-PtdIns(4,5)P_2_ (diC8PIP2), for two main reasons: i) an inositol phosphate binding site was described at the top of the C-lobe of both βarr1 and βarr2 (*30, 57*), and ii) the same analogue diC8PIP2 was shown to be important for NTSR1**-**βarr1 complex formation (*28*). Briefly, V2R was mixed with an equimolar concentration of diC8PIP2 (Avanti Polar lipids, Inc), an excess of βarr1ΔCT (2:1 molar ratio) and an excess of ScFv30 (2:1 molar ratio) as well as 250 µM AVP and 2.5 mM MgCl 2. In a representative experiment, concentrations of the different components of the complex were as follows: 35 µM V2R and diC8PIP_2_, 70 µM βarr1ΔCT and ScFv30. The coupling reaction was allowed to proceed at room temperature (RT) for 90 min. The complex was then purified through an orthogonal affinity chromatography procedure followed by a size exclusion chromatography step. First, to remove excess of βarr1ΔCT and ScFv30, the complex AVP-V2R-βarr1ΔCT-ScFv30 was purified by an M2 anti-Flag affinity chromatography. The mixture was loaded three times on the column, the resin was washed three times with 10 CV of wash buffer containing 20 mM HEPES (pH 7.5), 100 mM NaCl, 0.002 % CHS, 0.02 % LMNG, 0.005 % GDN, 10 µM AVP. The complex and the uncomplexed V2R were then eluted with 5 CV of wash buffer supplemented with Flag peptide (400 µg/ml). Second, the eluate was then loaded onto Strep-Tactin affinity resin to get rid of the uncomplexed receptors. The resin was washed with 10 CV of wash buffer (20 mM HEPES (pH 7.5), 100 mM NaCl, 0.02 % LMNG, 0.005 % GDN, 0.002 % CHS, and 10 µM AVP)). The complex was then eluted with 5 CV of wash buffer complemented with 2.5 mM desthiobiotin. Finally, the eluate was concentrated with a 50-kDa MWCO concentrator, and subjected to a SEC Superose 6 (10/300 GL, GE Healthcare) equilibrated with a buffer containing 20 mM Hepes (pH 7.5), 100 mM NaCl, 0.0011 % LMNG, 0.001 % GDN, 0.002 % CHS, and 10 µM AVP. The complex displayed a monodisperse peak whose analysis by SDS polyacrylamide gel and Coomassie blue staining confirmed the presence of all proteins (**Suppl Fig. S3**). Peak fractions were pooled, supplemented with amphipol A8-35 0.001 %, and concentrated using a 50-kDa MWCO concentrator to ∼3 mg/ml for cryo-EM studies.

#### Negative stain microscopy observations

Before preparing cryo-EM grids, we first checked the quality and the homogeneity of the AVP-V2R-βarr1ΔCT-ScFv30 samples by negative stain-electron microscopy (NS-EM). Three microliters (µl) of each complex at 0.04 mg/ml were applied for 2 min on glow-discharged carbon-coated grids and then negatively stained with uranyl formate 0.75 % for 1 min. Observation of EM grids was carried out on a JEOL 2200FS FEG Transmission Electron Microscope (TEM) operating at 200 kV under low-dose conditions (total dose of 20 electrons/Å^2^) in the zero–energy loss mode with a slit width of 20 eV. Images were recorded on a 4K × 4K slow-scan charge-coupled device camera (Gatan Inc.) at a nominal magnification of ×50,000 with defocus ranging from 0.5 to 1.5 µm. In total, 55 micrographs were recorded, allowing us to pick 97,182 particles using e2boxer from Eman2 package (*58*). Further processing was performed with Relion 3.1. The particles were subjected to a 2D classification including to get rid of free micelles and dissociated components of the complex. From 2D classes, 65,090 particles corresponding to the AVP-V2R-βarr1ΔCT-ScFv30 complexes were selected, representing 67% of all particles picked, a good prerequisite for cryo-EM analysis. The AVP­V2R-βarr1ΔCT-ScFv30 complex revealed a homogeneous distribution of particles showing a two-domain organization (V2R in the detergent micelle *versus* βarr1ΔCT-ScFv30). The two-dimensional (2D) class averages clearly showed the βarr1ΔCT tightly engaged within the micelle-embedded V2R (**Suppl Fig. S3**), an architecture in agreement with the “core” conformation of GPCR-βarr1 complex (*29*).

#### Cryo-EM sample preparation and image acquisition

For AVP-V2R-βarr1ΔCT-ScFv30 cryoEM investigation, 3 µl samples were applied on glow-discharged Quantifoil R1.2/1.3 300-mesh UltrAufoil grids (Quantifoil Micro Tools GmbH, Germany), blotted for 3.5 s, and then flash-frozen in liquid ethane using the semi-automated EM GP2 (Leica Microsystems) plunge freezer (100 % humidity and 4 °C). Images were collected in one session at the European Molecular Biology Laboratory (EMBL) of Heidelberg (Germany) on a FEI Titan Krios (Thermo Fisher Scientific) at 300 keV through a Gatan Quantum 967 LS energy filter using a 20 eV slit width in zero-loss mode and equipped with a K3 Summit (Gatan Inc.) direct electron detector configured in counting mode. Movies were recorded at a nominal energy-filtered transmission electron microscope magnification of ×130,000 corresponding to a 0.64 Å calibrated pixel size. The movies were collected in 40 frames in defocus range between −1 and −2 µm with a total dose of 52.63 *e*^−^/Å^2^. Data collection was fully automated using SerialEM, resulting in 14,080 movies.

#### Cryo-EM data processing

Movie frames were aligned and summed using the MotionCorr Relion own implementation (Relion 3.1.2) with 7 by 5 patches, a B-factor of 150 and a binning factor of 2, resulting in motion-corrected images with a pixel size of 1.28 Å. The contrast transfer function (CTF) parameters were estimated using Gctf (*59*). The images with a maximal resolution estimation worse than 7 Å were discarded, resulting in 13,566 images (**Suppl Fig. S5**). A first automatic picking was carried out using boxnet from Warp software package (*60*), allowing to select and extract 3,610,370 particles which were transferred into Relion v3.1.2. Iterative 2D classifications combined with bad 2D class averages exclusion sorted out a total of 1,169,437 particles. Those particle coordinates were used as references to train a model with Topaz (*61*), a positive-unlabeled convolutional neural network for particle picking. It resulted in the picking of 4,595,394 particles. Those particles were transferred into Relion 3.1.2 and subjected to iterative 2D classifications. The best resulting particles obtained from boxnet (1,806,545 particles) and Topaz (2,660,410 particles) were then merged and duplicate removed yielding 3,721,020 particles. Iterative 2D classifications yielded a total of 729,335 projections from best 2D class averages (**Suppl Fig. S4**). Several rounds of CryoSPARC v3.2.0 were performed using 2D classification and *ab initio* reconstruction processing steps (6 times using two models) resulting in the selection of a particle stack comprising 27,637 particles which yielded to a density map with an overall resolution of 4.75 Å after three-dimensional non-uniform (NU) refinement (**Suppl Fig. S6**). Resolution were estimated using the gold-standard FSC at 0.143 (FSC = 0.143). The final particle stack was transferred into Relion for micelle-V2R signal subtraction. The particle boxes were recentered and adapted to the size of the βArr1-ScFv30 complex. Subtracted particles were subjected to local refinement in CryoSPARC yielding to a density map (EMD-14223) with an overall resolution of 4.23 Å (FSC = 0.143) (**Suppl Fig. S7**). Attempts to align the complex with micelle subtraction yielded density maps with similar apparent quality and overall resolution than without subtraction (r ∼ 4.8-5 Å). Attempts to align the V2R alone were unsuccessful. The subset of 27,637 particles was then refined using additional *ab-initio* steps to obtain a subset of 8,296 particles which yielded to a density map (EMD-14221) with an overall resolution of 4.73 Å after NU refinement (**Suppl Fig. S7**). This map displays more detailed features of the dynamic regions of the complex as compared to the initial 27,637-particle density map. Local resolution of the two maps calculated using CryoSPARC (EMD-14221 and EMD-14223) ranged from 3.5 to 5.5 Å.

#### Model building and refinement

For each of the two cryo-EM maps - EMD-14221 (full-complex) and EMD-14223 (focused around the βArr1ΔCT-ScFv30 substructure) - an atomic model was built and refined. A starting model of V2RCter-βarr1ΔCT-ScFv30 substructure was built using the atomic structure of the V2RCter (residues 356 to 368 according to residue numbering of UniProt entry P30518) from PDB-6u1n (4 Å resolution), and the Fab30 and βarr1 from PDB-4jqi (2.6Å resolution). The model was fitted into the EMD-14223 map and manually adjusted (sequence mutations and loop reconstruction) using Coot (*62*). Then, this intermediate model was rigid-body fitted into the EMD-14221 map and completed with the structure of the V2R TM domain+AVP from PDB­7kh0 (2.8Å resolution) and a molecule of diC8PIP2. The resulting model of the full complex (AVP-V2R-βarr1ΔCT-ScFv30) was real-space refined against the EMD-14221 map under standard stereochemical restraints (including Ramachandran restraints) using Coot (*62*) and Phenix (*63*). The refined model of the full complex (PDB-7Rr0c) does not include the V2R N-ter portion (residues 1-31), parts of the V2R intracellular loops (ICLs) (residues 148-156 from ICL1; residues 183-188 in ICL2; residues 239-263 in ICL3) and parts of the V2R Cter (residues 343 to 355 and residues 369-371) which were not visible in the density maps. The model of βarr1ΔCT in PDB-7r0c includes residues 6 to 365 whereas ScFv30 is nearly complete (residues 110-128 are missing). The intermediate model of the V2RCter-βarr1ΔCT-ScFv30 substructure (initially fitted in the EMD-14223 map) was further real-space refined (with Phenix and Coot) against the EMD-14223 map by temporarily including the V2R TM domain from PDB-7r0c (in order to avoid βarr1ΔCT residues jumping into V2R residual density still present in proximity of βarr1ΔCT) and by imposing extremely strong geometrical constraints on the V2R model (in order to avoid unjustified modifications of the V2R regions farther from the βarr1ΔCT where no density was left after the signal subtraction procedure). The V2R TM domain was finally removed from the refined V2RCter-βarr1ΔCT-ScFv30 model (PDB-7r0j).

#### Liquid chromatography – tandem mass spectrometry (LC-MS/MS) and analysis of V2R phosphoresidues

To prepare samples for LC-MS/MS, the purification of the V2R was adapted from a previously described protocol (see above, paragraph V2R expression and purification) (*22*). As indicated, the receptor was first purified by affinity chromatography using Strep-Tactin resin (IBA) following the same protocol. Then, the eluate was then loaded onto a M2 anti-Flag affinity resin (Sigma-Aldrich). The column was washed first with 10 CV of a buffer containing 20 mM HEPES (pH7.5), 100 mM NaCl, 0.1 % (w/v) DDM, 0.01 % (w/v) CHS, and 10 µM TVP. A second wash followed with 10 CV of a buffer containing 20 mM HEPES (pH 7.5), 100 mM NaCl, 0.025 % (w/v) DDM, 0.005 % (w/v) CHS, and 10 µM TVP. The bound receptor was eluted in the same buffer supplemented with FLAG peptide (0.4 mg/ml) (5 CV). The fractions corresponding to the receptor were collected and the HRV3C protease was added for overnight cleavage at 4 °C (at a 1:20 (HRV3C:V2R) weight ratio). After concentration using a 50-kDa MWCO concentrator (Millipore), the V2R was purified by SEC using a Superdex 200 increase (10/300 GL column) connected to an ÄKTA purifier system (GE Healthcare). Fractions corresponding to the pure monomeric receptor were pooled (∼1.5 ml) and concentrated to 30 µM (1.2 mg/ml). The purified V2R (100 to 200 µg) was digested using micro S-Trap columns (https://protifi.com/, Huntington NY) following the supplier’s protocol. Briefly, after reduction (20 mM DTT 10 min 95 °C) and alkylation (40 mM IAA 30 min in the dark), the receptor was digested using 3 µg of trypsin (Promega, Gold) for 1 hour at 47 ° C. The peptides obtained were analysed using nano-throughput HPLC (Ultimate 3000-RSLC, Thermo Scientific) coupled to a mass spectrometer (Qexactive-HF, Thermo Scientific) equipped with a nanospray source. The pre-concentration of the samples was carried out in-line on a pre-column (0.3 mm × 10 mm, Pepmap®, Thermo Scientific) and separation of the peptides on a column (0.075 mm × 500 mm, reverse phase C18, Pepmap®, Dionex) following a gradient from 2 to 25 % buffer B (0.1 % AF in 80 % ACN) for 100 min at a flow rate 300 nl / min, then 25 to 40 % in 20 min and finally 40 to 90 % in 3 minutes. The spectra were acquired in mode: “data-dependent acquisition” (dynamic exclusion of 20 seconds). The LC-MS / MS analysis cycle is therefore composed of several phases, a “Full scan MS” with analysis in the orbitrap at 60,000 resolution followed by MS/MS (HCD fragmentation), for the 12 most abundant precursors at a resolution of 30,000. Raw spectra were processed using the MaxQuant environment v1.6.10.43 (*64*) and Andromeda for database search with match between runs and the iBAQ algorithm enabled. The MS/MS spectra were matched against the sequence of the V2R construct (**Suppl Fig. S11**), the reference proteome (Proteome ID UP000008292, release 2021_01) of Autographa californica nuclear polyhedrosis virus, the UniProt entries (release 2021_01, https://www.uniprot.org/) for Spodoptera frugiperda (Fall armyworm, taxon identifier 7108) and 250 frequently observed contaminants, as well as reversed sequences of all entries. Enzyme specificity was set to trypsin/P, and the search included cysteine carbamidomethylation as a fixed modification and oxidation of methionine, acetylation (protein N-term) and phosphorylation of Ser, Thr, Tyr residue (STY) as variable modifications. Up to two missed cleavages were allowed for protease digestion. The maximum false discovery rate for peptides and proteins was set to 0.01. Representative protein ID in each protein group was automatically selected using the in-house developed Leading tool v3.4 (*65*). Signal intensities of receptor peptides were extracted using Skyline v2.1.0.31(*66*), with the option « Use high-selectivity extraction ». For a defined peptide sequence, a score corresponding to the probability of phosphorylation for each possible position (S, T or Y) was determined (**Suppl Fig. S11**). The normalized sum of all these probabilities is then used to define the confidence of localization, known as localization probability (*67*). Classically, class I phosphosites correspond to sites with a localization probability of at least 0.75 (*68*).

#### Time-resolved Fluorescence Resonance Energy Transfer binding assays

V2R binding studies using TagLite assays (PerkinElmer Cisbio) based on time-resolved fluorescence resonance energy transfer (TR-FRET) measurements were previously described (*22, 48, 69*). Briefly, HEK cells were plated in white-walled, flat-bottom, 96-well plates (Greiner CELLSTAR plate, Sigma-Aldrich) in Dulbecco’s minimum essential medium (DMEM) containing 10% fetal bovine serum (Eurobio), 1% nonessential amino acids (GIBCO), and penicillin/streptomycin (GIBCO) at 15,000 cells per well. Cells were transfected 24 hours later with a plasmid coding for the V2R version used in cryo-EM studies fused at its N-terminus to the SNAP-tag (SNAP-V2R) (PerkinElmer Cisbio). Transfections were performed with X-tremeGENE 360 (Merck), according to the manufacturer’s recommendations: 10 µl of a premix containing DMEM, X-tremeGENE 360 (0.3 µl per well), SNAP-V2 coding plasmid (30 ng per well), and noncoding plasmid (70 ng per well) were added to the culture medium. After a 48-hour culture period, cells were rinsed once with Tag-lite medium (PerkinElmer Cisbio) and incubated in the presence of Tag-lite medium containing 100 nM benzylguanine-Lumi4-Tb for at least 60 min at 37 °C. Cells were then washed four times. For saturation studies, cells were incubated for at least 4 hours at 4 °C in the presence of benzazepine-red nonpeptide vasopressin antagonist (BZ-DY647, PerkinElmer Cisbio) at various concentrations ranging from 1 × 10^−10^ to 1 × 10^−7^ M. Nonspecific binding was determined in the presence of 10 µM vasopressin. For competition studies, cells were incubated for at least 4 hours at 4 °C with benzazepine-red ligand (5 nM) and increasing concentrations of vasopressin ranging from 1 × 10^−11^ to 3.16 × 10^−6^ M. Fluorescent signals were measured at 620 nm (fluorescence of the donor) and at 665 nM (FRET signal) on a PHERAstar (BMG LABTECH). Results were expressed as the 665/620 ratio [10,000 × (665/620)]. A specific variation of the FRET ratio was plotted as a function of benzazepine-red concentration (saturation experiments) or competitor concentration (competition experiment). All binding data were analyzed with GraphPad 9.1.1 (GraphPad Prism Software Inc.) using the one site-specific binding equation. All results are expressed as the means ± SEM of at least three independent experiments performed in triplicate (**Suppl Fig. S2**). *K*_i_ values were calculated from median inhibitory concentration values with the Cheng-Prusoff equation.

#### cAMP accumulation assays

V2R functional studies based on TR-FRET measurements were described previously (*22, 51, 70*). Briefly, HEK cells were plated in black-walled 96-well plates (Falcon) at 15,000 cells per well. Cells were transfected 24 hours later with a plasmid coding for the V2R version used in cryo-EM studies. Transfections were performed with X-tremeGENE 360 (Merck), according to the manufacturer’s recommendations: 10 µl of a premix containing DMEM, X-tremeGENE 360 (0.3 µl per well), SNAP-V2 coding plasmid (0.002 ng per well), and noncoding plasmid (100 ng per well) were added to the culture medium. After a 24-hour culture period, cells were treated for 30 min at 37 °C in the cAMP buffer with or without increasing AVP concentrations (3.16 × 10^−12^ to 10^−6^ M) in the presence of 0.1 mM RO201724, a phosphodiesterase inhibitor (Sigma-Aldrich). The accumulated cAMP was quantified using the cAMP Dynamic 2 Kit (PerkinElmer Cisbio) according to the manufacturer’s protocol. Fluorescent signals were measured at 620 and 665 nm on a Spark 20M multimode microplate reader (Tecan). Data were plotted as the FRET ratio [10,000 × (665/620)] as a function of AVP concentration [log (AVP)]. Data were analyzed with GraphPad Prism v 9.1.1 (GraphPad Prism Software Inc.) using the “dose-response stimulation” subroutine. Median effective concentrations were determined using the log(agonist) versus response variable slope (four parameters) fit procedure. Experiments were repeated at least three times on different cultures, each condition in triplicate. Data are presented as means ± SEM (**Suppl Fig. S2**).

#### β-arrestin recruitment assays

Upon GPCR activation, β-arrestins (βArr) are recruited to stop G protein signaling and to initiate clathrin-mediated receptor internalization. During this process, the release of the C-terminal domain of βArrs is associated with the binding of βArrs to the adaptor protein 2 (AP2). This interaction can be measured using the HTRF® technology (PerkinElmer CisBio) based on the use of two specific antibodies, one directed against β-arrestin2 (βArr2), the second one specific for AP2. In this assay (βArr2 recruitment kit, PerkinElmer CisBio), the AP2 antibody is labeled with a Europium (Eu) cryptate fluorescent donor, and the one against βArr2 is labeled with a d2 fluorescent acceptor, their proximity being detected by FRET signals. The specific signal is positively modulated in proportion with the recruitment of βArr2 to AP2 upon V2R activation by AVP. Briefly, HEK cells were plated at a seeding density of 2.5×10^4^ cells per well in a white-walled 96-well plates (CELLSTAR plate, Sigma-Aldrich) precoated with poly-L-ornithine (14 µg/ml, Sigma-Aldrich) for 24 hours, in Dulbecco’s minimum essential medium (DMEM) (GIBCO) complemented with 10 % fetal bovine serum (FBS) (Eurobio), 1 % non-essential amino acids (GIBCO), and 1 % penicillin-streptomycin antibiotics solution (GIBCO). To produce the V2R, the cells were transfected with 30 ng of the pRK5-Flag-Snap-V2R plasmid (coding for the cleaved V2R construct used in cryo-EM studies) using X-trem gene 360 (Merck), according to the manufacturer’s recommendations. After a 24-hour culture, cells were used to evaluate the recruitment of βArr2 to AP2 upon V2R activation with the βArr2 recruitment kit (PerkinElmer CisBio) following the manufacturer recommendation. Briefly, the cells were first washed one time with DMEM-free and incubated 2 hours at RT with 100 ul per well of stimulation buffer containing various concentrations of the ligand AVP (ranging from 10^−6^ M to 10^−12^ M). The media was then replaced by 30 µl per well of stabilization buffer for 15 min at RT. The cells were then washed three times with 100 µl per well of wash buffer before adding 100 µl per well of a pre-mix of Eu cryptate and d2 antibodies in detection buffer. Following overnight incubation at RT, 80 ul of media were removed from each well before reading the 96-well plates on a Spark 20M multimode microplate reader (Tecan) or a PHERAstar FS (BMG Labtech) by measuring the signals of the donor (Europium cryptate-labeled AP2 antibody) at a wavelength of 620 nm, and of the acceptor at 665 nm (d2-labeled βArr2). Finally, the results were expressed as the FRET ratio [(665/620) x 10,000] and plotted using GraphPad 9.1.1 (GraphPad Prism software inc.). Experiments were repeated at least three times on different cultures, each condition in triplicate. Data are presented as means ± SEM (**Suppl Fig. S2**).

### Molecular Dynamic (MD) simulation method

MD simulations were performed for the AVP-V2R-βarr1ΔCT complex with and without diC8PIP2 (**Suppl Fig. S9 and S10**). The initial models were built from the cryo-EM structure reported here. The missing ICL2 was modeled based on the cryoEM structure of AVP-V2R-Gs (PDB 7dw9) (*24*). Other missing loop regions were generated by Modeller v9.15 (*71*). Residues S241^ICL3^, T253^ICL3^, S255^ICL3^, S357^Cter^, T359^Cter^, T360^Cter^, S362^Cter^, S363^Cter^ and S364^Cter^ of V2R were phosphorylated. PACKMOL-Memgen (*72*) was used to assign the side-chain protonation states and embed the models in a bilayer of POPC lipids. and the system was solvated in a periodic 120 × 120 × 140 Å^3^ box of explicit water and neutralized with 0.15 M of Na^+^ and Cl^­^ions. We used the Amber ff14SB-ildn (*73*) and lipid 14 (*74*) force fields, as well as the Amber force field parameters for phosphorylated amino acids (*75*). The TIP3P (*76*) and the Joung-Cheatham (*77*) parameters were used for the water and the ions, respectively. Effective point charges of the phosphoinositide were obtained by RESP fitting (*78*) of the electrostatic potentials calculated with the HF/6-31G* basis set. After energy minimization, all-atom MD simulations were carried out using Gromacs 5.1 (*79*) patched with the PLUMED 2.3 plugin (*80*). Each system was gradually heated to 310 K and pre-equilibrated during 10 ns of brute-force MD in the *NPT-*ensemble. The replica exchange with solute scaling (REST2) (*81*) technique was used to enhance the sampling of the loop regions at the V2R-arrestin interface. A total of 24 replicas were simulated in the *NVT* ensemble. REST2 is a type of Hamiltonian replica exchange simulation scheme, which performs many replicas of the same MD simulation system simultaneously. The replicas have modified free energy surfaces, in which the barriers are easier to cross than in the original system. By frequently swapping the replicas during the MD, the simulations “travel” on different free energy surfaces and easily visit different conformational zones. Finally, only the samples on the original free energy surface are collected. The replicas are artificial and are only used to overcome the energy barriers. REST2, in particular, modifies the free energy surfaces by scaling (reducing) the force constants of the “solute” molecules in the simulation system. In this case, the loop regions at the V2R-arrestin interface were considered as “solute”–the force constants of their van der Waals, electrostatic and dihedral terms were subject to scaling–in order to facilitate their conformational changes. The effective temperatures used here for generating the REST2 scaling factors ranged from 310 K to 1000 K, following a distribution calculated with the Patriksson-van der Spoel approach (*82*). Exchange between replicas was attempted every 1000 simulation steps. This setup resulted in an average exchange probability of ∼40 %. We performed 120 ns × 24 replicas of MD in the *NVT* ensemble for each system. The first 40 ns were discarded for equilibration. The original unscaled replica (at 310 K effective temperature) was collected and analyzed.

---

**Fig. S1.**
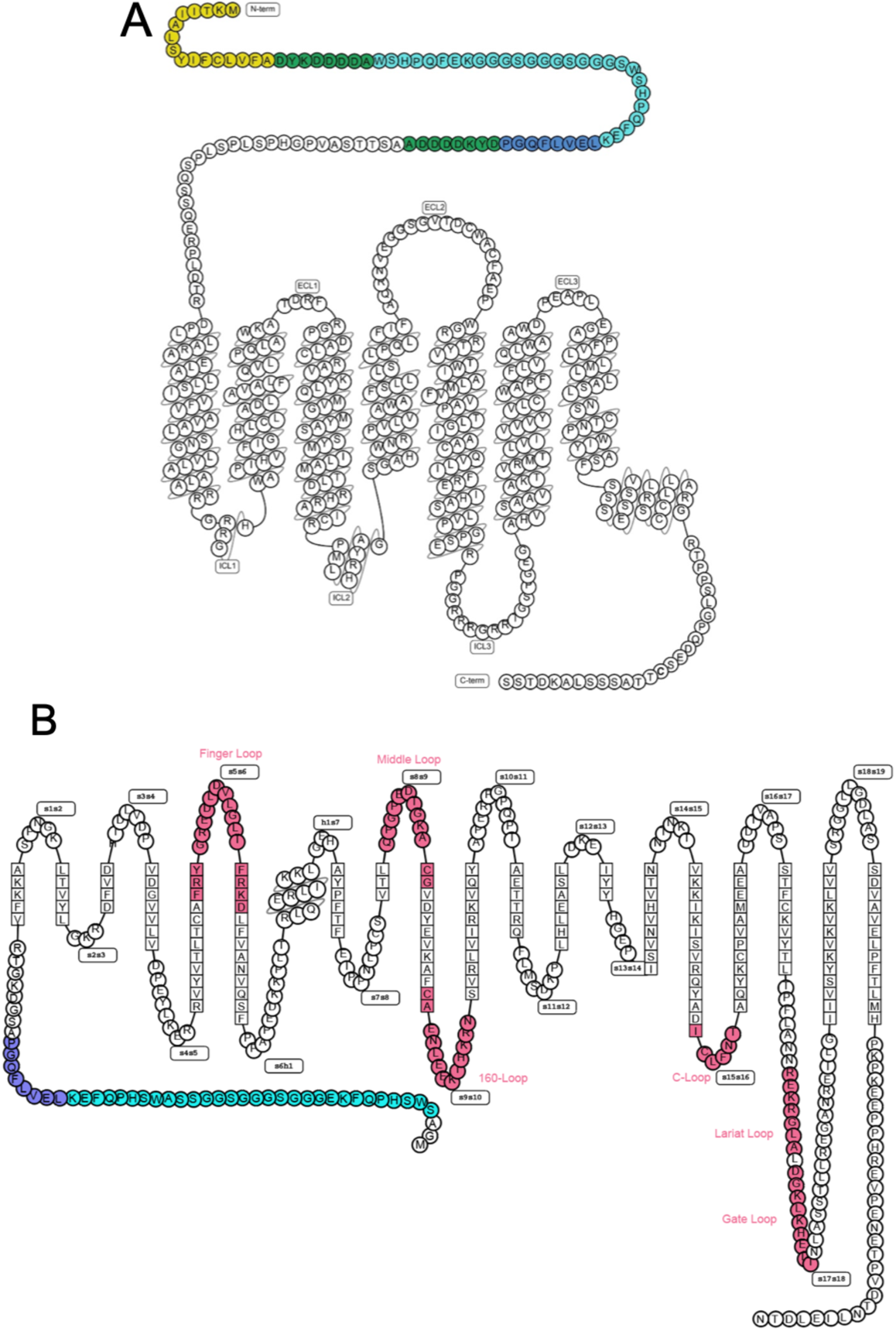
Cartoons of the V2R and βarr1ΔCT constructs. (**A**) and (**B**) Modified snake plots of the engineered V2R and βarr1 versions used for Cryo-EM structure determination, respectively (https://gpcrdb.org). Purification tags were inserted in the N-terminal part of both constructs for different reasons: i) the C-terminus of V2R has to be maintained in a native form for an efficient βarr1 interaction, ii) the C-terminus of βarr1 was truncated at residue 382 (βarr1ΔCT), leading to a constitutively active form enhancing V2R coupling. The hemagglutinin signal peptide is shown in yellow, the Flag-tags are in green, the Twin-Strep-tags are in cyan, the human rhinovirus 3C protease cleavage site are in purpleblue, and all βarr1ΔCT loops discussed in the manuscript are depicted in pink.

**Fig. S2.**
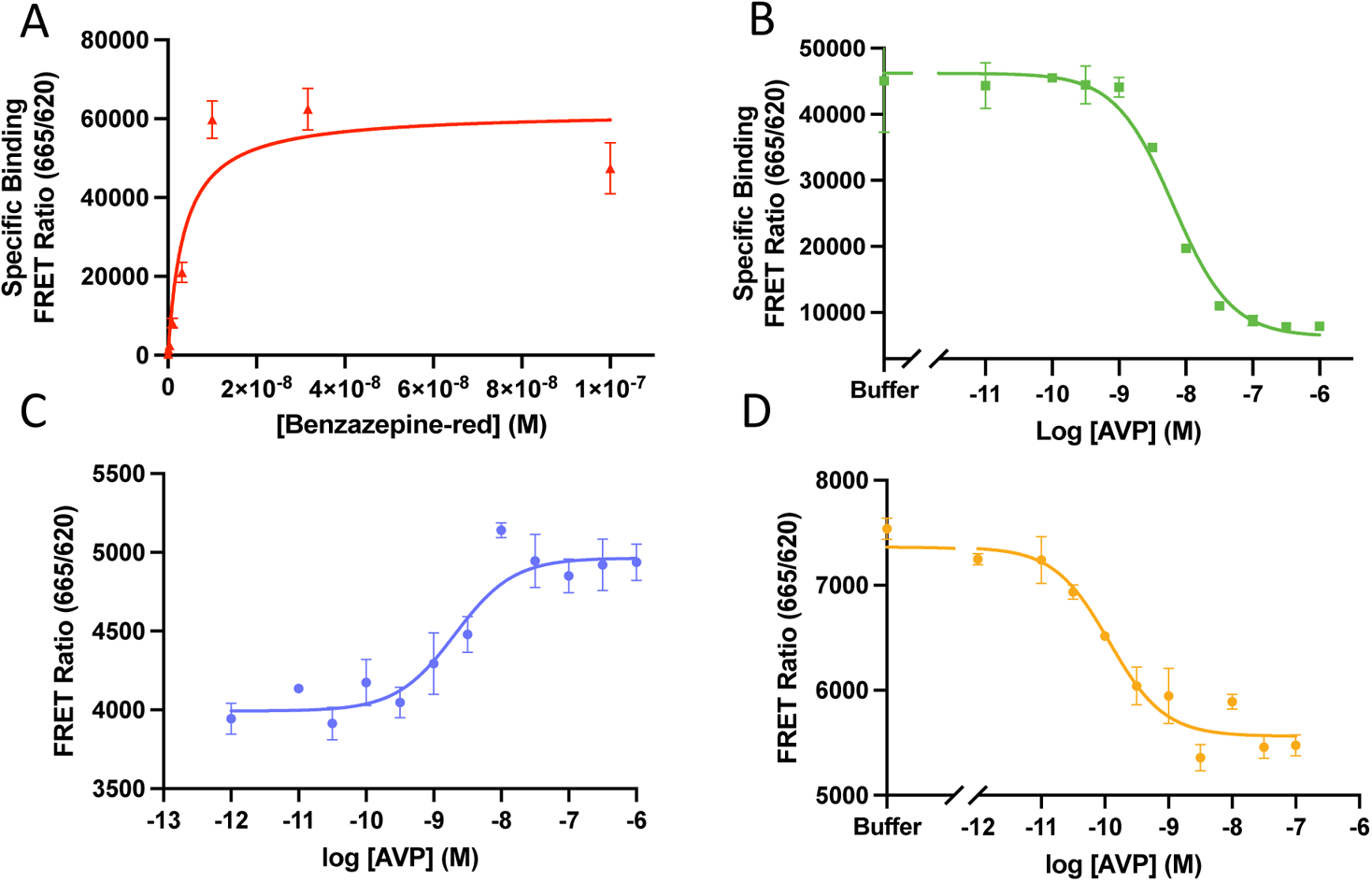
Pharmacological and functional properties of the V2R construct. (**A**) Binding of the benzazepine-red fluorescent antagonist to the V2R measured by FRET. Specific binding from a typical saturation assay is shown as FRET ratio (665nm/620nm x 10,000). (**B**) Binding of AVP to the V2R is illustrated as FRET ratio (665nm/620nm x 10,000). Specific binding of the fluorescent antagonist is shown. For each competition curve, it was used at 5 nM with or without increasing concentrations of AVP. (**C**) Dose-response of V2R-dependent recruitment of the βarr2 to AP2 measured by FRET ratio (665nm/620nm x 10,000) in the presence of increasing concentrations of AVP. (**D**) Dose-response of V2R-dependent Gs protein/adenylyl cyclase activation measured by FRET ratio (665nm/620nm x 10,000). The cAMP accumulation which displaces the fluorescently-labeled cAMP binding to its specific antibody is shown in the presence of increasing concentrations of AVP. For each assay, a typical experiment is represented from at least 3 independent experiments, each point performed in triplicate. Each value is presented as mean ± SEM.

**Fig. S3.**
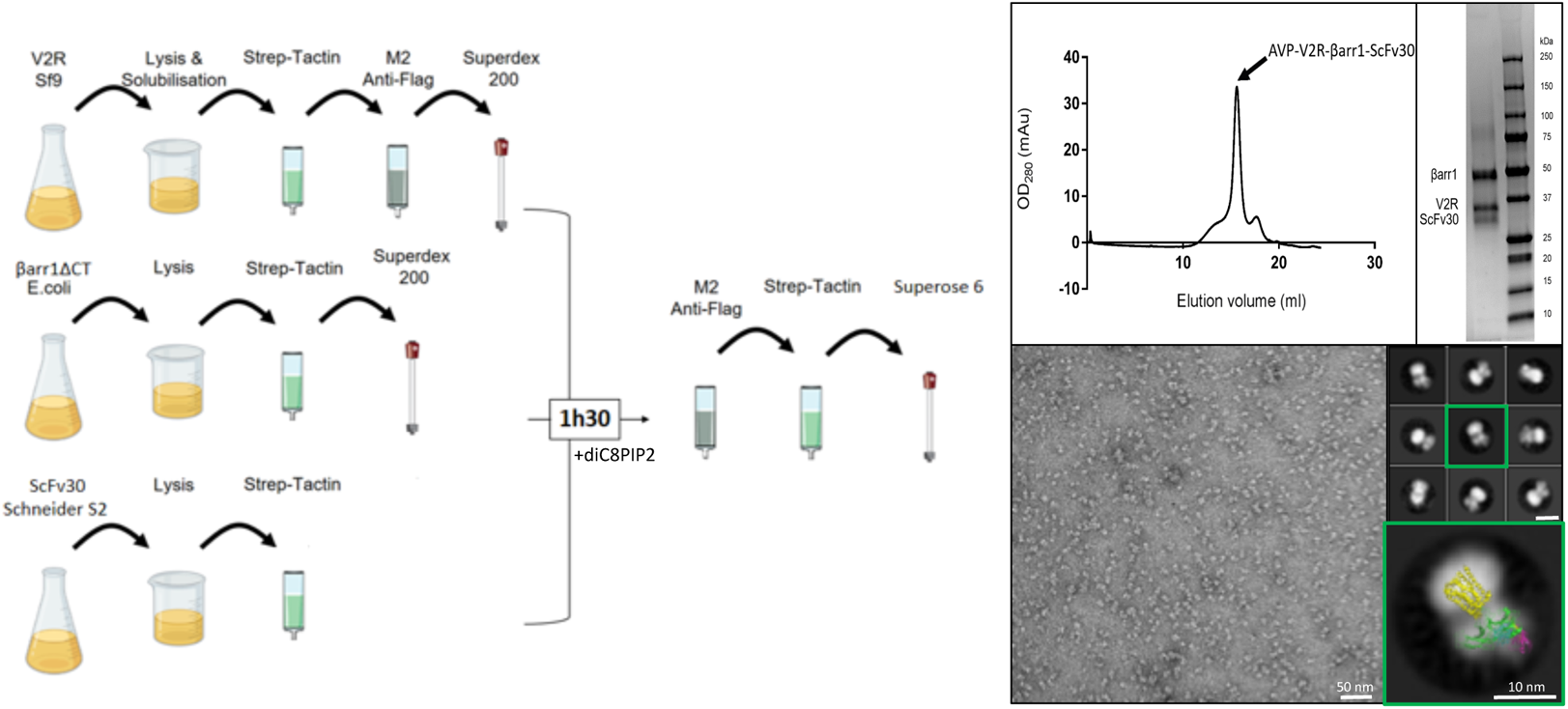
Overview of the AVP-V2R-βarr1ΔCT-ScFv30 complex preparation, purification and NS-EM analysis. (**A**) Workflow for AVP-V2R-βarr1ΔCT-ScFv30 assembly. The V2R, βarr1ΔCT and ScFv30 were expressed and purified separately, the complex being incubated for 1h30 in the presence of diC8PIP2 and then isolated by two successive affinity chromatographies and a final SEC step. (**B**) A representative chromatogram of the AVP-V2R-βarr1ΔCT-ScFv30 complex using a Superose 6 column shows a monodisperse peak. Fractions containing the sample were combined, used directly for NS-EM and concentrated for Cryo-EM grid preparation. SDS-PAGE of peak fraction from the Superose 6 step is shown on the right. Coomassie blue staining of the proteins confirmed the presence of βarr1ΔCT, V2R and ScFv30 in the complex (AVP and diC8PIP2 are not visible). (**C**) NS-EM analysis of the sample. A representative micrograph of the purified AVP-V2R-βarr1ΔCT-ScFv30-diC8PIP2 complex isolated from the SEC peak (scale bar, 50 nm) is shown on the left. 2D most representative class averages showing different orientations (scale bar, 10 nm) are illustrated (top right). A close-up view of a typical 2D class average is presented (bottom right). The 3D model of the M2R-βarr1-Nb24-ScFv30 complex (PDB 6u1n) is fitted onto the particle to show that the AVP-V2R-βarr1ΔCT-ScFv30 particle displayed typical size and characteristics of a GPCR-βarr1-ScFv30 assembly (M2R, yellow; βarr1, green; Nb24, blue; ScFv30, pink).

**Fig. S4.**
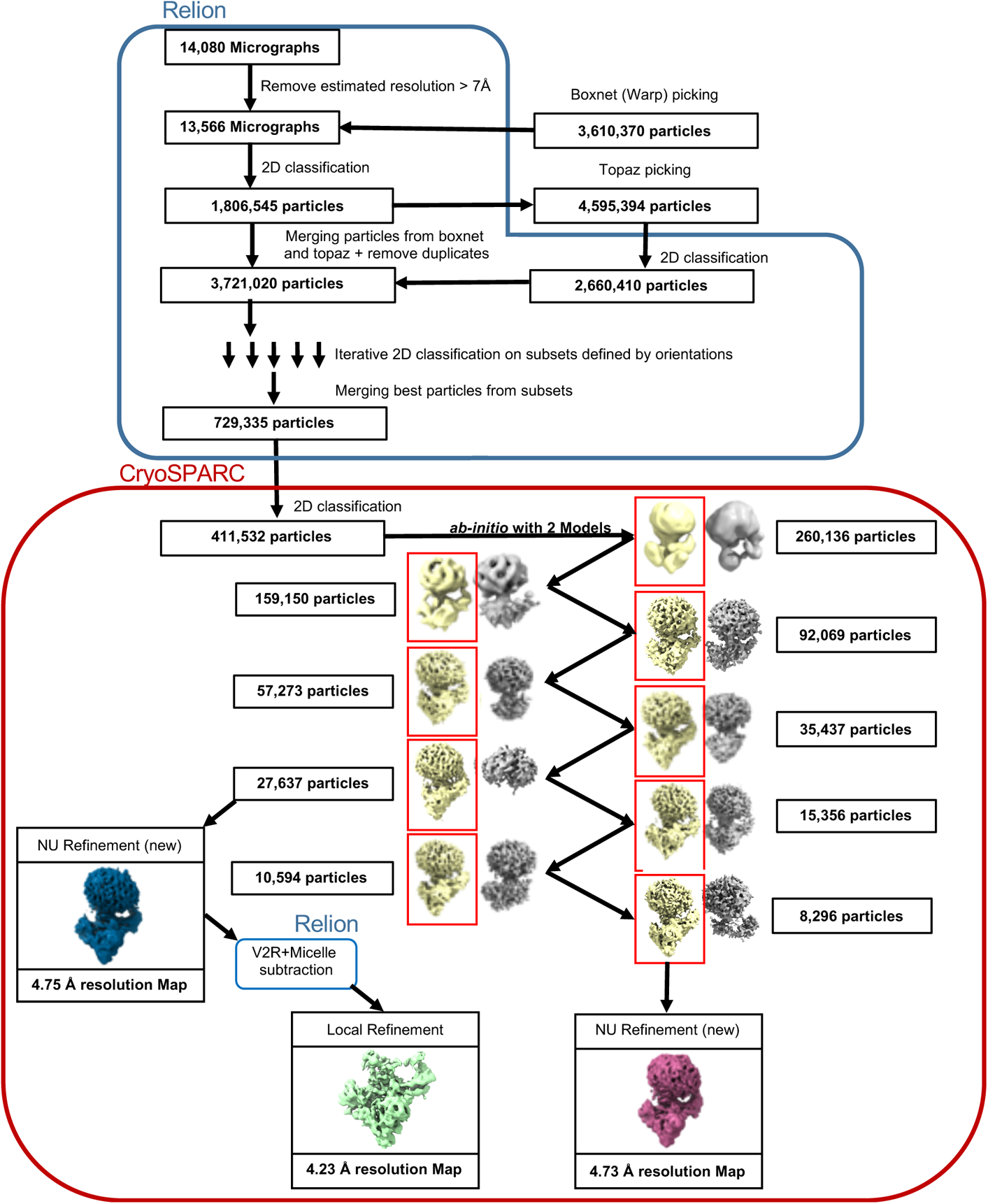
Cryo-EM workflow. The different steps of the single particle analysis from a unique movie dataset collected with a Titan Krios are detailed. Micrographs were treated to screen particles which were subjected to iterative 2D classification in Relion. Using this 2D classification and ab initio reconstructing processing steps in CryoSPARC, a stack of 27,637 particles was selected which generated a density map with an overall resolution of 4.75 Å after non uniform refinement. Subtraction of micelles and V2R and local refinement yielded another map with an overall resolution of 4.23 Å. In addition, the subset of 27,637 particles was also refined using three additional ab-initio steps to obtain another stack of 8,296 particles yielding a density map with an overall resolution of 4.73 Å after NU refinement.

**Fig. S5.**
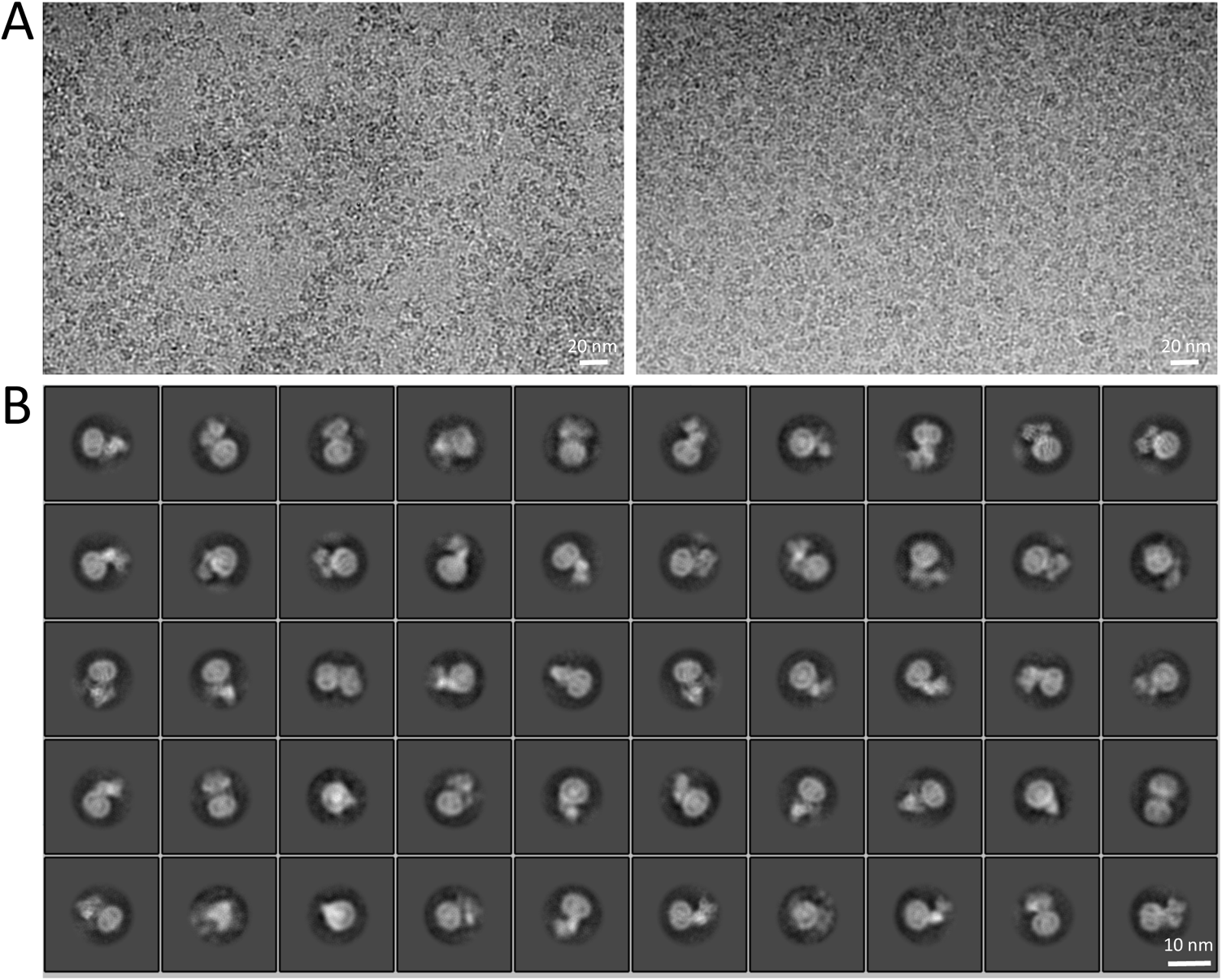
Cryo-EM images and 2D class averages of the AVP-V2R-βarr1ΔCT-ScFv30 complex. (**A**) Two representative micrographs of the complex, showing a different distribution of particles in ice. (**B**) Most representative 2D class averages showing distinct secondary structure features like the V2R TM regions embedded in the detergent micelle, and different orientations of the AVP-V2R-βarr1ΔCT-ScFv30 complex.

**Fig. S6.**
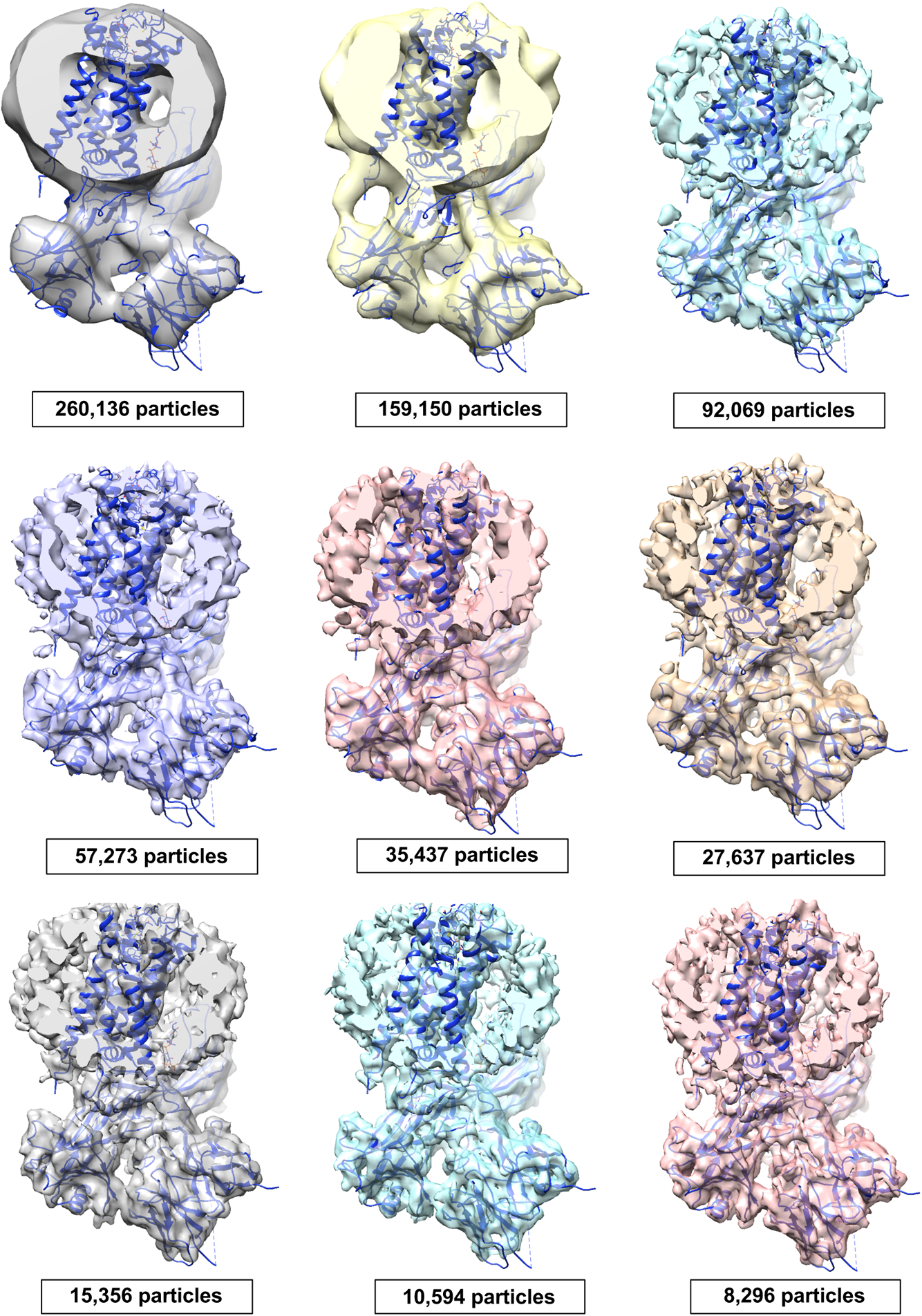
Screening of the particles with a 2 models ab initio reconstruction procedure. The refined 3D model fitted in the successive density maps shows that orientation of βarr1ΔCT-ScFv30 relative to V2R is maintained through selected models from the iterative rounds of CryoSPARC *ab initio* analysis.

**Fig. S7.**
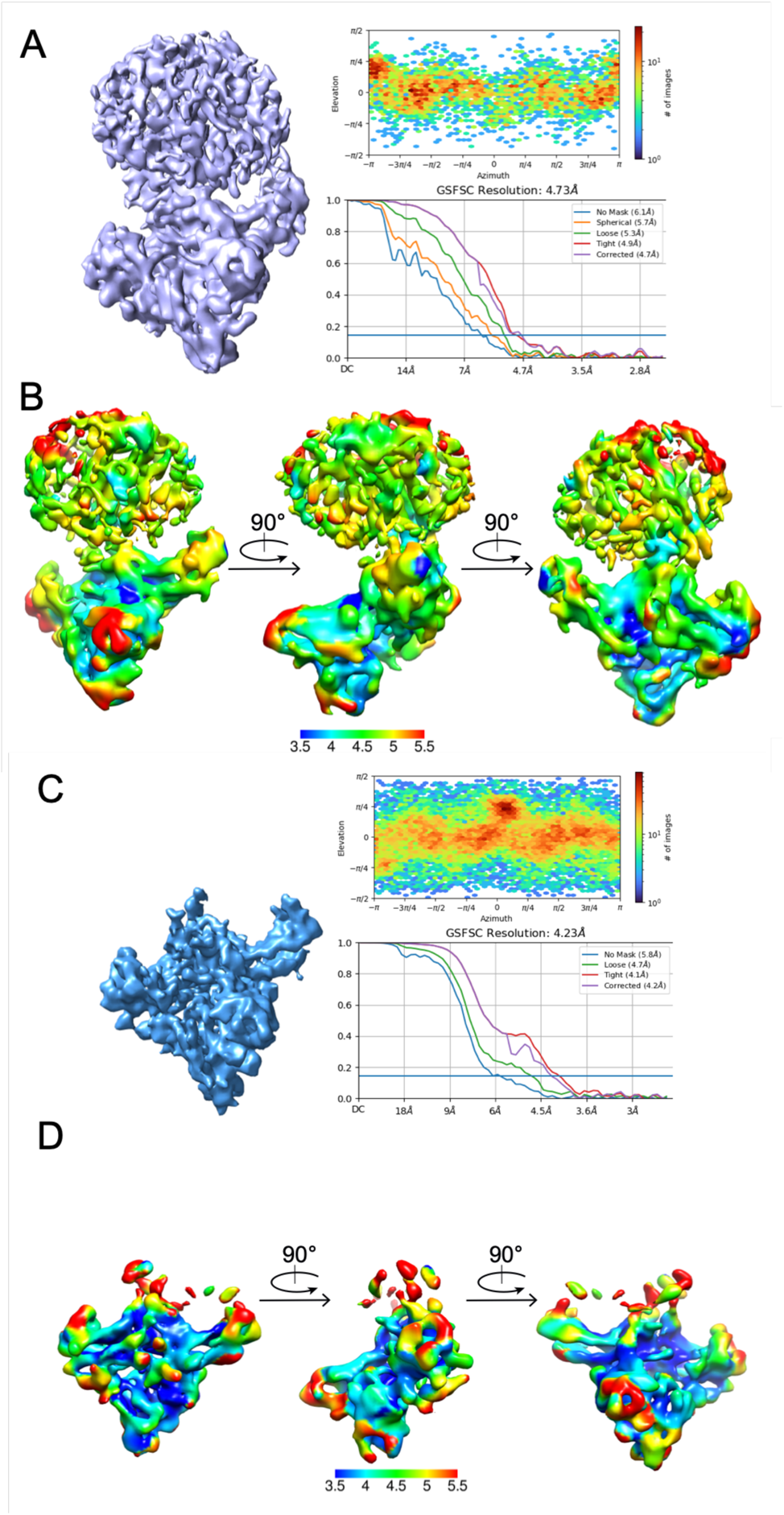
Cryo-EM density maps of the AVP-V2R-βarr1ΔCT-ScFv30 complex and local resolution estimation. (**A**) Final density map of the whole AVP-V2R-βarr1ΔCT-ScFv30 complex and (**B**) its local resolution estimation. (**C**) Final density map of the V2RCter-βarr1ΔCT-ScFv30 subcomplex and (**D**) its local resolution estimation. For both complexes, viewing direction distribution (top panels) indicates that no preferential orientation of the particles was observed. The density map resolution was determined from FSC with a cut-off of 0.143 (bottom panel).

**Fig. S8.**
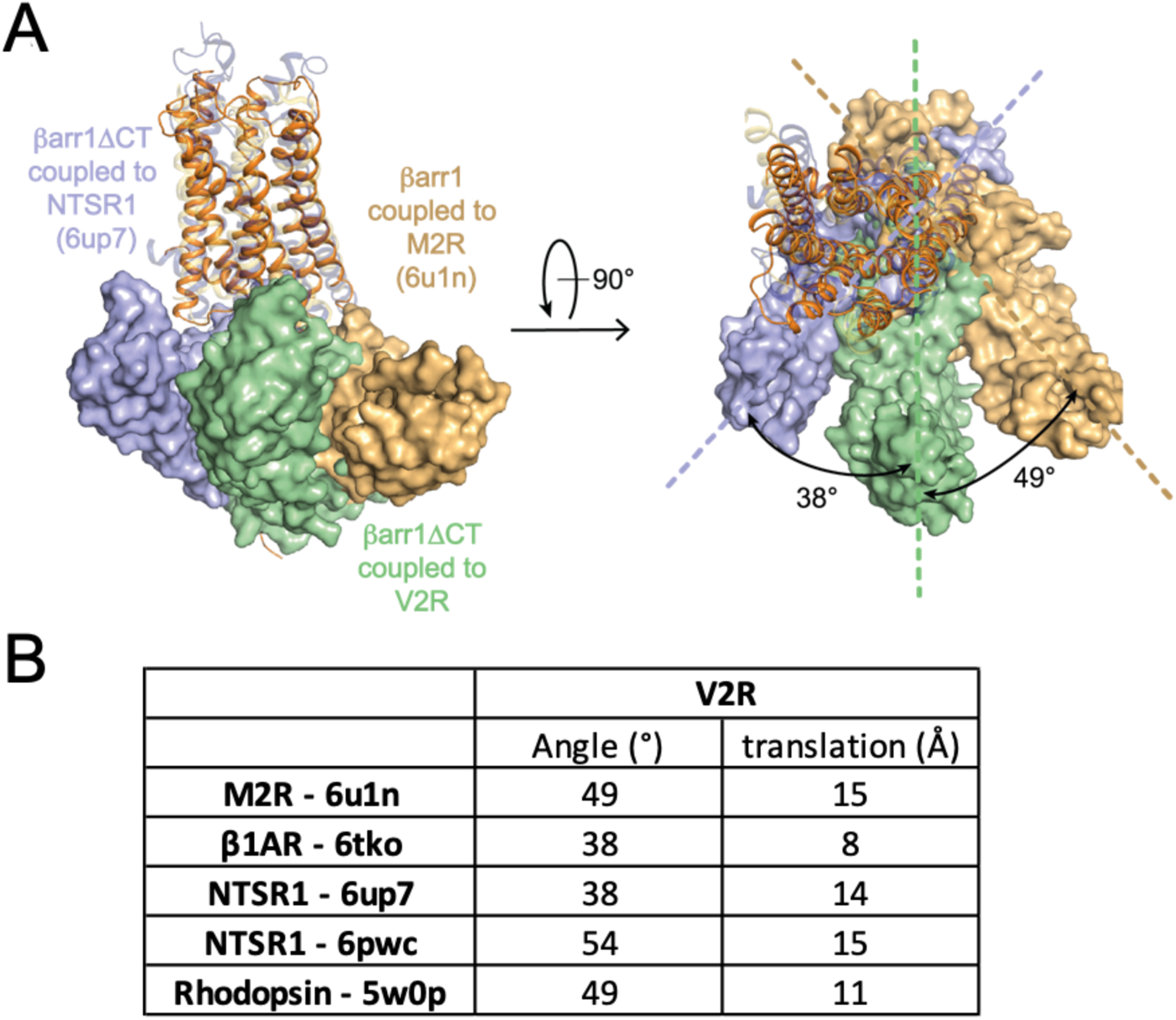
Comparison of the βarr1ΔCT orientation in different GPCR complexes. (**A**) Overlay of the V2R-βarr1ΔCT structure with NTSR1-βarr1ΔCT (6up7) and M2R-βarr1 (6u1n) structures, on the basis of alignment of the receptor chains, viewed from the membrane (left) and from the extracellular space (right). The orientation of βarr1ΔCT in the V2R complex is compared with that in the NTSR1 and in the M2R complexes respectively, and the angles between βarr1s are indicated. V2R and βarr1ΔCT are colored in orange and green, M2R and βarr1 (6u1n) are in yellow and gold, βarr1ΔCT and NTSR1 (6up7) are in purple. (**B**) Geometrical parameters of the different GPCR-arrestin complexes compared to the V2R-βarr1ΔCT complex. The rhodopsin-arrestin1 (PDB 5w0p) is also indicated for information (*83*). Angles and distances were extracted from Pymol 2.3.5 macro’s “angle_between_domains”. They represent rotation and displacement that would happen to align βarr1s after each couple of GPCRs is aligned.

**Fig. S9.**
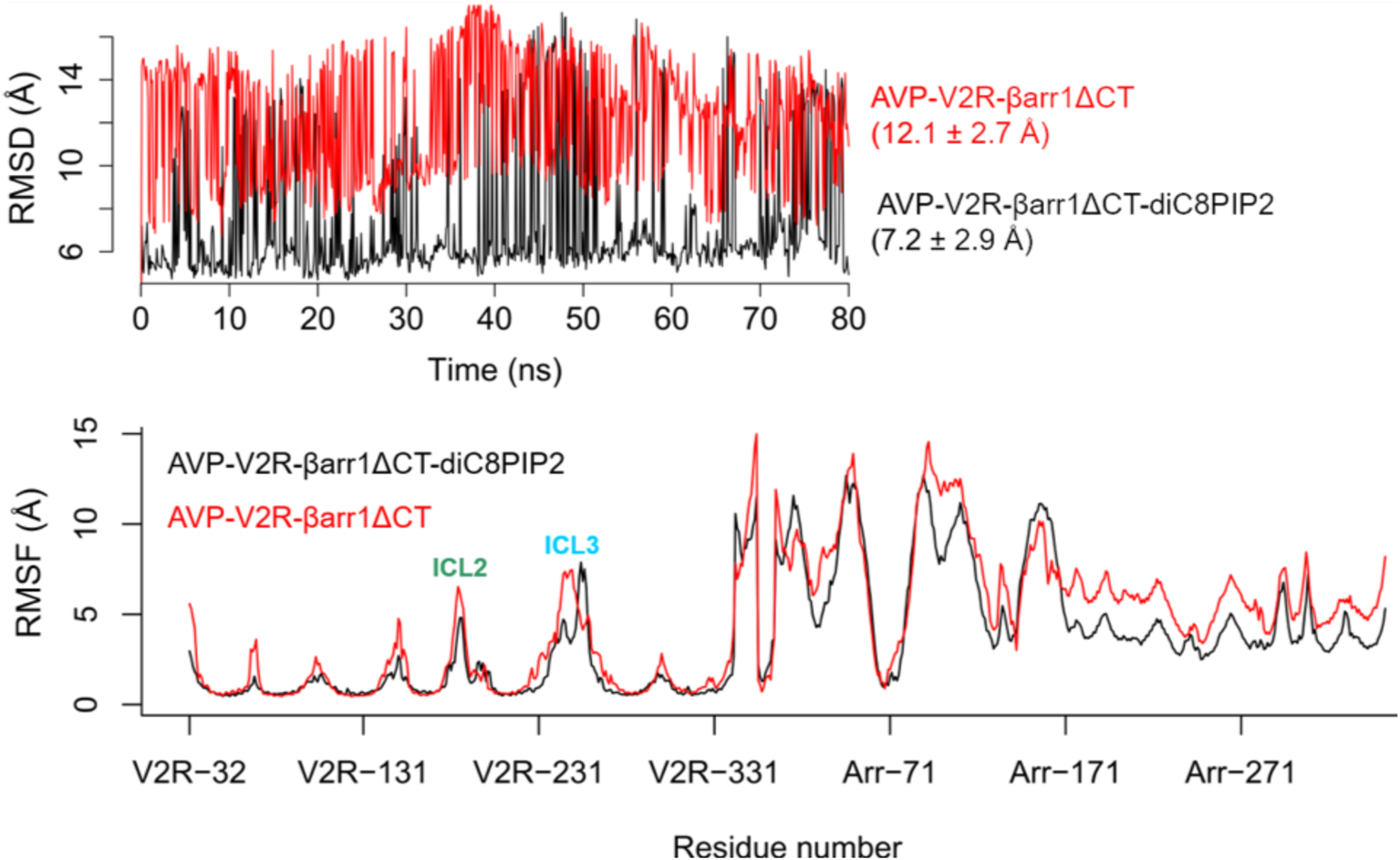
MD simulations of the AVP-V2R-βarr1ΔCT complex: effect of diC8PIP2. Root-mean-square deviations (RMSD, top panel) and fluctuations (RMSF, bottom panel) of V2R-βarr1ΔCT Cα atoms, during the MD simulations of the AVP-V2R-βarr1ΔCT complex with or without diC8PIP2. The MD trajectories were aligned to the Cα atoms of V2R TM helices in the cryo-EM structure.

**Fig. S10.**
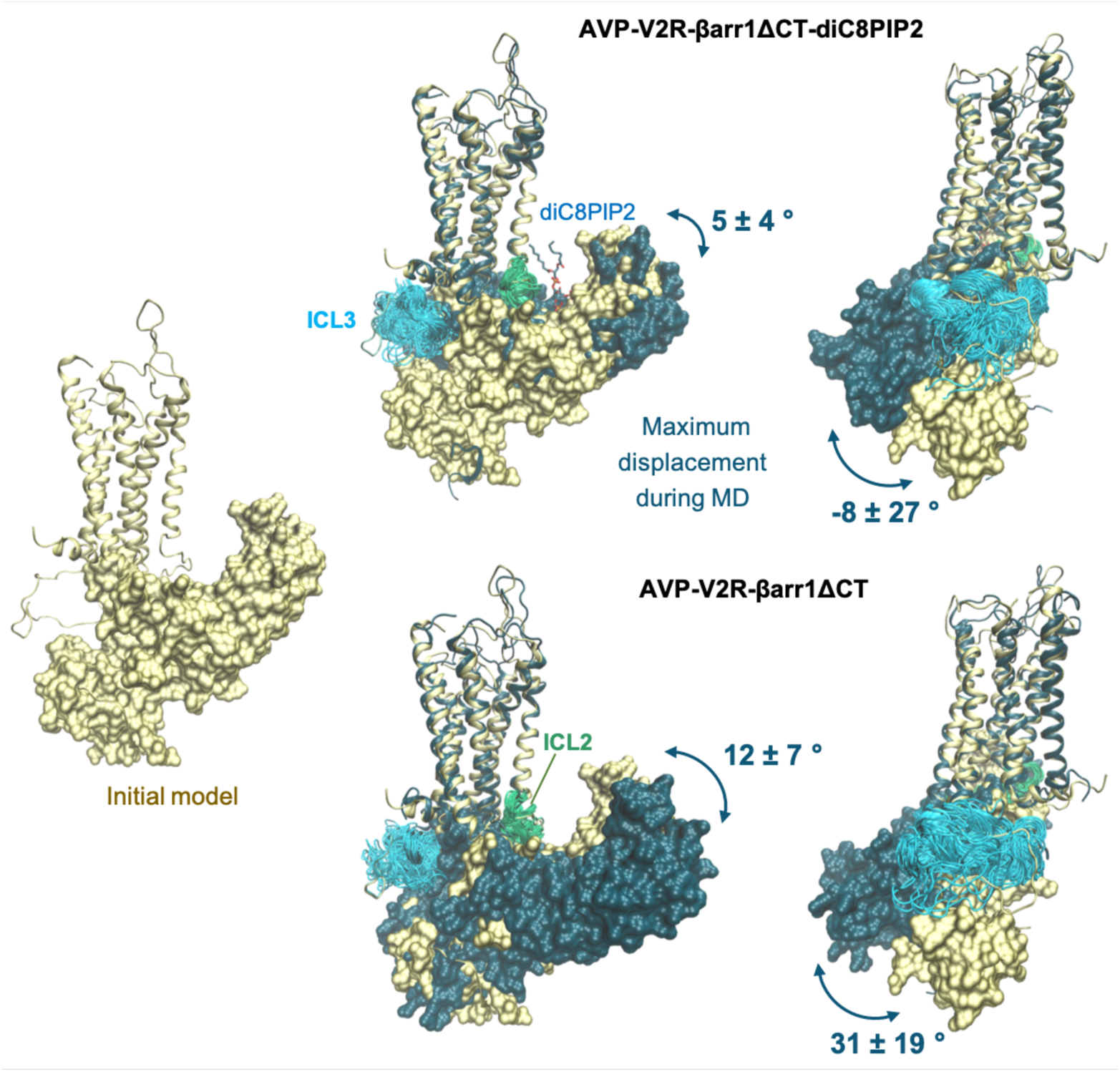
MD simulations of the AVP-V2R-βarr1ΔCT complex: mobility of V2R ICLs and βarr1ΔCT. The mobility of V2R ICL2, ICL3 and βArr1ΔCT during the MD simulations of the AVP-V2R-βArr1ΔCT complex with (top) or without (bottom) diC8PIP2 is shown. MD trajectories were aligned to the Cα atoms of V2R TM helices in the initial model built based on the Cryo-EM structure. Rotation of βArr1ΔCT N-lobe is measured as a dihedral angle with respect to the axis of TM2. Tilt of βArr1ΔCT C-lobe is calculated as a dihedral angle with respect to the membrane plane. All angles are calculated relative to the initial model, labeled as mean ± SD during the MD.

**Fig. S11.**
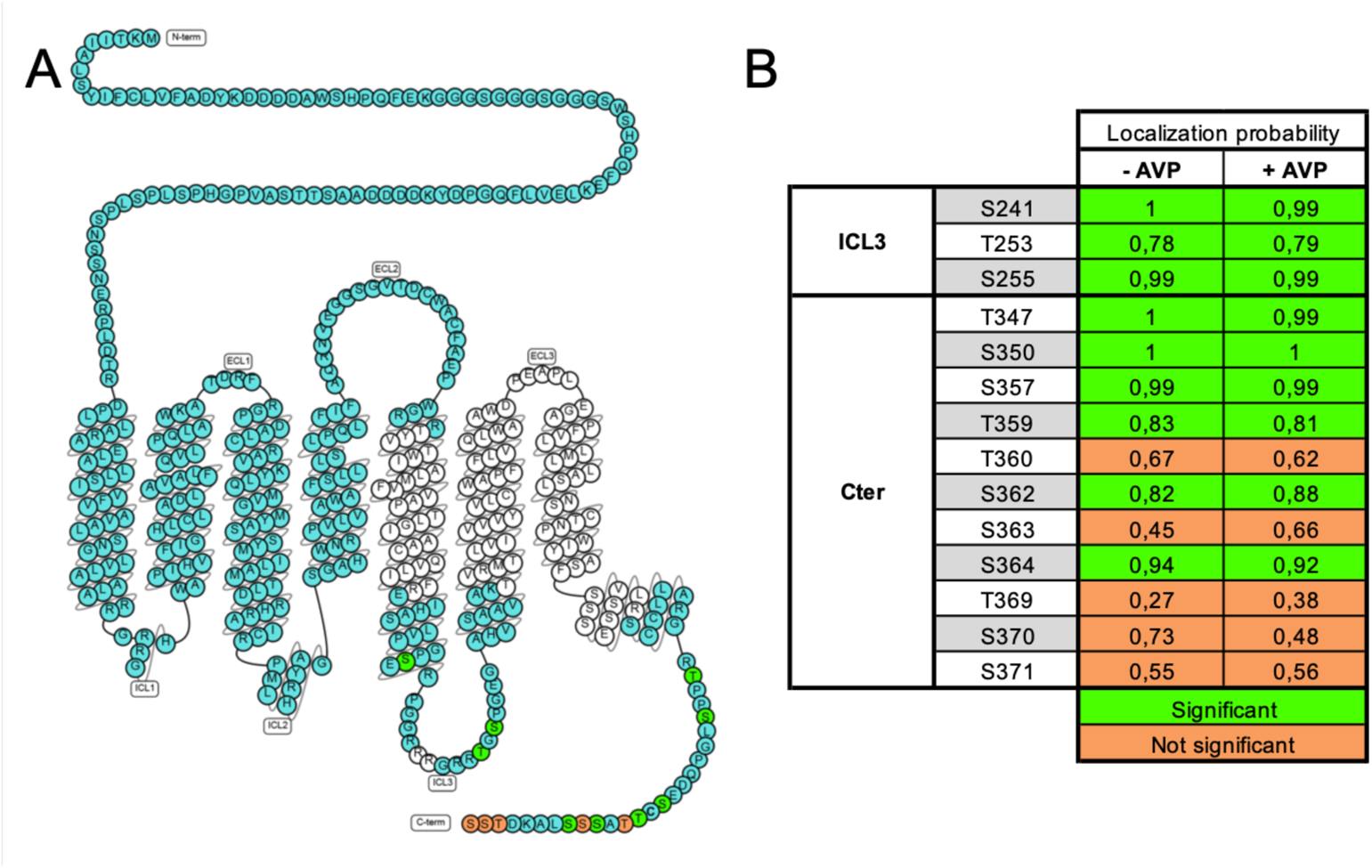
Phosphoproteomics of the Cryo-EM V2R version. (**A**) Modified snake plot of the engineered V2R used for cryo-EM structure determination. Suspensions of Sf9 cells expressing the recombinant V2R were treated or not with AVP 1 μM for 30 min at 28°C before harvesting. The V2R was then purified using the procedure described in Materials and Methods, isolated as a monodisperse peak by SEC and subjected to trypsin digestion. Peptides were analyzed using nano-HPLC and tandem mass spectrometry. Identified peptides allowed to recover most (approximately 70%) of the V2R sequence (shown in cyan). Phosphoresidues identified by LC-MS/MS are in green and orange. (**B**) Localization probability of phosphoresidues. Phosphosites were identified in ICL3 and V2RCter regions from cells treated by AVP or not. Significant localization probability (at least 0.75) of phosphates is shown in green, whereas the not significant (< 0.75) is shown in orange.

**Table S1.** Cryo-EM data collection, refinement, and validation statistics. EMDB, Electron Microscopy Data Bank, PDB, Protein Data Bank, RMSD, root mean square deviation.

|  | AVP-V2R-βArr1ΔCT-ScFv30<br>(PDB 7R0C; EMD-14221) | V2RCter-βArr1ΔCT-ScFv30<br>(PDB 7R0J; EMD-14223) |
| --- | --- | --- |
| **Data collection and processing** |  |  |
| Voltage (kV) | 300 | 300 |
| Electron exposure (e-/Å<sup>2</sup>) | 52.63 | 52.63 |
| Defocus range (μm) | -1.0 to -2.0 | -1.0 to -2.0 |
| Pixel size (Å) | 0.64 | 0.64 |
| Symmetry imposed | C1 | C1 |
| Initial particle images (no.) | 3,721,020 | 3,721,020 |
| Final particle images (no.) | 8,293 | 27,637 |
| Map resolution (Å) | 4.73 | 4.35 |
| FSC threshold | 0.143 | 0.143 |
| Map local resolution range (Å) | 3.5 to 5.5 | 3.5 to 5.5 |
| **Refinement** |  |  |
| Initial models used (PDB codes) | 7KH0, 4JQI, 6U1N | 4JQI, 6U1N |
| Model resolution (Å) | 4.73 | 4.23 |
| Map sharpening *B* factor (Å<sup>2</sup>) | -168.2 | -188 |
| Model composition |  |  |
| Non-hydrogen atoms | 6,905 | 4,713 |
| Number of protein residues / atoms | 884 / 6,865 | 603 / 4,713 |
| Number of ligands / ligand atoms | 1 / 40 | 0 / 0 |
| average *B* factor (Å<sup>2</sup>) |  |  |
| Protein | 224 | 165 |
| Ligands | 254 | — |
| R.m.s deviations |  |  |
| Bond lengths (Å) | 0.005 | 0.007 |
| Bond angles (°) | 1.11 | 1.16 |
| Validation |  |  |
| MolProbity score | 2.32 | 2.57 |
| Clashscore | 25 | 37 |
| Poor rotamers (%) | 0.14 | 0.19 |
| Ramachandran plot |  |  |
| Favored (%) | 93.81 | 91.30 |
| Allowed (%) | 6.19 | 8.7 |
| Disallowed (%) | 0 | 0 |

**Table S2.** Contact map between V2R ICL3 and βArr1ΔCT during MD simulations. Proposed contacts between residues of V2R ICL3 (row) and βArr1ΔCT (column) by MD simulations of the AVP-V2R-βArr1ΔCT complex in the presence or absence of diC8PIP2. They are colored by the lifetime during the simulations (% of the total simulation frames). A 6 Å minimum distance cutoff between residue pairs was used to calculate the contacts in each MD simulation frame. For instance, in the presence of diC8PIP2, R247 from V2R and G132 from βArr1ΔCT formed the most frequent contact (in 87% of the simulation frames).

|  | V40 | V41 | L42 | E50 | R51 | R52 | V53 | Y54 | V55 | T56 | P124 | C125 | S126 | V127 | T128 | L129 | Q130 | P131 | G132 | P133 | E134 | D135 | T136 | G137 | K138 |
| --- | --- | --- | --- | --- | --- | --- | --- | --- | --- | --- | --- | --- | --- | --- | --- | --- | --- | --- | --- | --- | --- | --- | --- | --- | --- |
| AVP-V2R-βarr1ΔCT |  |  |  |  |  |  |  |  |  |  |  |  |  |  |  |  |  |  |  |  |  |  |  |  |  |
| G239 |  | 5 |  |  |  |  |  |  |  |  |  |  |  |  |  |  |  |  |  |  |  |  |  |  |  |
| S241 |  | 13 | 1 |  |  |  |  |  |  |  |  |  |  |  |  |  |  |  |  |  |  |  |  |  |  |
| E242 |  |  |  |  |  |  |  |  |  |  |  |  |  |  |  |  |  |  |  |  |  |  |  |  |  |
| R243 |  |  | 7 |  |  |  |  |  |  |  |  |  |  |  |  |  |  |  |  |  |  |  |  |  |  |
| P244 |  | 3 | 9 |  |  |  |  |  |  |  |  |  |  |  |  |  |  |  |  |  |  |  |  |  |  |
| G245 |  | 6 | 1 |  |  |  |  |  |  |  |  |  |  | 3 |  |  |  |  |  |  |  |  |  |  |  |
| G246 |  | 9 | 13 |  |  |  |  |  |  |  |  | 3 | 15 | 19 | 8 |  | 1 |  | 3 | 1 |  | 3 | 4 | 1 | 2 |
| R247 | 5 | 18 | 26 |  |  |  |  | 3 | 7 | 1 | 13 | 1 | 4 | 23 | 9 | 22 | 36 | 9 | 16 | 7 | 3 | 7 | 5 | 3 | 2 |
| R248 |  |  | 7 |  |  |  |  |  |  |  |  |  |  | 1 | 2 | 1 | 1 |  |  |  | 1 | 1 | 1 |  |  |
| R249 |  |  | 3 |  | 6 | 2 | 4 | 11 | 14 | 3 |  |  | 2 | 16 | 3 | 1 | 5 |  | 4 |  |  | 1 | 1 |  |  |
| G250 |  |  |  | 5 | 8 | 3 | 7 | 17 | 17 | 6 |  |  | 1 | 7 | 3 | 2 | 16 | 2 | 16 |  | 1 | 1 |  |  |  |
| R251 |  |  |  |  | 3 |  | 1 | 7 | 9 | 1 |  |  | 3 | 5 | 6 | 6 |  |  |  |  |  |  |  |  |  |
| R252 |  | 1 | 2 |  | 1 |  |  | 8 | 1 | 1 |  |  |  |  |  |  |  |  |  |  |  |  |  |  |  |
| T253 |  |  | 1 |  |  |  |  |  |  |  |  |  |  |  |  |  |  |  |  |  |  |  |  |  |  |
| P256 |  |  | 2 |  |  |  |  |  |  |  |  |  |  |  |  |  |  |  |  |  |  |  |  |  |  |
| AVP-V2R-βarr1ΔCT-diC8PIP2 |  |  |  |  |  |  |  |  |  |  |  |  |  |  |  |  |  |  |  |  |  |  |  |  |  |
| G246 |  |  |  |  |  |  |  |  |  |  |  |  |  |  |  |  |  |  |  |  |  |  |  |  |  |
| R247 |  | 11 | 2 |  |  |  |  | 6 | 2 |  |  |  |  | 2 | 3 | 4 | 19 | 74 | 87 | 33 | 36 |  |  |  |  |
| R248 |  |  |  |  |  |  |  |  |  |  |  |  |  | 2 |  | 2 | 6 | 10 | 15 | 4 | 10 |  |  |  |  |
| R249 |  |  |  |  |  |  |  |  |  |  |  |  |  | 4 | 4 | 4 | 9 | 16 | 50 | 6 | 31 |  |  |  |  |
| G250 |  |  |  |  |  |  |  | 7 | 11 |  |  |  |  | 6 | 3 | 3 | 7 | 1 | 4 | 6 | 18 |  |  |  |  |
| R251 |  |  |  |  |  |  |  | 8 | 7 | 2 |  |  |  |  |  |  |  |  |  |  | 9 |  |  |  |  |
| R252 |  |  |  |  |  |  |  |  |  |  |  |  |  | 2 | 1 | 1 |  |  | 1 |  | 9 |  |  |  |  |
| T253 |  |  |  |  |  |  |  |  |  |  |  |  |  |  |  |  |  |  |  |  | 3 |  |  |  |  |

